# Comparative genomic analysis reveals novel phylogenetically intermediate Streptococci with high phenotypic diversity in the human distal lung microbiota

**DOI:** 10.1101/2023.12.22.572891

**Authors:** Slipa Kanungo, Germán Bonilla-Rosso, Garance Sarton-Lohéac, Marianne Kuffer, Markus Hilty, Thomas Geiser, Philipp Engel, Sudip Das

**Author notes:** Philipp Engel and Sudip Das jointly supervised the work. #Address correspondence to Sudip Das. Present address: Slipa Kanungo, Instituto Gulbenkian de Ciência, Oeiras, Portugal. Germán-Bonilla Rosso, Bioinformatics and Proteogenomics, Agroscope, Zürich, Switzerland.

## Abstract

Streptococci are one of the predominant and the most diverse genus in the human lung. Previously, we isolated human distal lung Streptococci from bronchoalvolear lavage fluid (BALF) as part of the human Lung Microbiota culture Collection (LuMiCol). Here, we performed whole genome sequencing, comparative phylogenomics and phenotypic characterization of six Streptococcal isolates representing the phylogenetic diversity of the genus in distal human lung. Here, we report five new species and one new subspecies including phylogenetic intermediates of commonly found Streptococci not limited to human lung. Pangenome analysis reveals gene content, evolutionary relationships, and metabolic functions shedding light on contribution of these Streptococci to lung microbial metabolism. Antimicrobial resistance gene analysis followed by MIC determination revealed macrolide, lincosamide and tetracycline resistance in lung Streptococci. We show the presence of capsular genes in lung streptococci both matching to the prototypical capsular genes (*cps*) and unique genes. Interestingly, the new *Streptococcus* isolate sp. nov. P2E5, genetically identical to the most prevalent *Streptococcus* in the human distal lung was revealed to be a phylogenetic intermediate between the *S. mitis* group and *S. pneumoniae.* It also harbors the pneumolysin (*ply*) gene and was found to have the serotype 21E. Finally, core genome phylogeny reveals that lung Streptococci the are evolutionary distinct from oral Streptococcal isolates in expanded Human Oral Microbiome Database (eHOMD). Hence, these findings we reveal new phylogenetically distinct Streptococcal species from the human distal lung microbiota and its genetic diversity and metabolism to understand the microbial ecology of human lung.

**Importance:** A healthy human distal lung harbour characteristic microbial communities mostly composed of oropharyngeal taxa, which are facultative or obligative anaerobes despite lung being the medium of oxygen intake. However, little is known about the genetic and functional diversity of these bacteria owing to the lack of resources including availability of primary lung isolate from human samples. Therefore, we have established a large bacterial collection that covers all major phyla by cultivating human bronchoalveolar lavage fluid (BALF) under various conditions. *Streptococcus* is the most prevalent and diverse genera in the human lung microbiota. Using genetic and biochemical approaches, we studied six diverse lung isolates from our collection representing the actual Streptococcal diversity and identify these as new species and subspecies. We hypothesize that learning about the phylogenetic genetic diversity, preferred metabolism and molecular structures of these Streptococci will provide with new insights on the understudied microbial ecosystem of the human lung.

## Introduction

The development of culture independent high-throughput DNA sequencing (both marker-gene amplicon sequencing and shotgun metagenomics) has made it possible to study the composition, diversity, and function of human microbial communities. Using culture-dependent and independent techniques we and others have shown that healthy human lungs harbour characteristic microbial communities(1–5). The lung microbiota is a complex and dynamic ecosystem composed of a diverse community of microorganisms, including bacteria, viruses, and fungi. The most prevalent bacterial phyla in lung are *Bacteroidetes* and *Firmicutes*, with low numbers of *Proteobacteria* and *Actinobacteria*(6). In addition, the biomass in the lung is relatively low compared to gut content with different genus level composition(7, 8). Most lung bacterial commensals have been shown to be of oral or supraglottic in origin(9, 10). However, the structure and composition of the lung bacterial communities are distinct(11). These differences may occur due to different oxygen conditions, pressure, pH, nutrients and distinct immune cell populations like airway macrophages that bacteria encounter during colonization(12). This suggests that the microbial ecology of lung and the interaction with the immune system is distinct from other sites on the human body(7, 12). Despite the implications of lung-associated bacteria in lung health(13–16), our current understanding of resident lung microbiota is poor. (2) With the advent of next generation sequencing came the ease of having taxonomic snapshot of a particular microbial niche leading to bacterial culture being overlooked. However, this is changing now with a broader realization of the importance of microbial cultivation and genotyping (17). In line with this, we performed large-scale culturing efforts with human BALF samples to obtain more than 300 bacterial isolates from 47 species to build the open source bacterial biobank called [LuMiCol (Lung Microbiota culture Collection](https://github.com/sudu87/Microbial-ecology-of-the-transplanted-human-lung), Figure S1A). This covers the most prevalent species in human lung as well as important pathogens, observed by amplicon sequencing. This is an important resource that will facilitate experimental work on the human lung microbiota. We have also demonstrated that *Streptococci* were the most prevalent and diverse genus within the balanced pneumotype supporting homeostasis(18). Streptococci were amongst the most prevalent OTUs (5 out of 22, 16S rRNA gene identity) in bronchoalveolar lavage fluids, which was also apparent when cultivated to establish our bacterial collection. Our culture collection harbored six representative isolates, five matched to the most prevalent and abundant Streptococci (OTU_11: P2E5, OTU_34: P2D11, OTU_42: P3D4, OTU_57: P3B4, 369.3: OTU_69) in human lung within our cohort (>97% 16S rRNA gene identity). Additionally, one isolate represented a rare *Streptococcus*.

Due to this diversity, we hypothesize that these Streptococci may represent the major metabolic pathways and provide valuable insights into the microbial ecology of the human distal lung. In this study, we sequenced the genomes of six *Streptococcus* isolates that represent each phylotype (97% 16S rRNA identity) followed by whole genome phylogenetic analysis, biochemical and metabolic tests. By doing these, we identified five new species and one novel subspecies of *Streptococcus*. Next, by employing pangenome analysis, we reveal orthologous gene content. Using a custom pipeline, we predicted metabolic pathways and macromolecular structures. We also reveal antibiotic resistance and virulence factors in these commensal streptococci. Finally, we compared lung streptococcal isolates with closely related genomes from expanded human oral microbiome database (eHOMD) to reveal their phylogenetic relationships.

## Results

### Whole genome phylogenomics and phenotypic characterisation identifies new streptococcal species and subspecies from distal human lung

We performed whole genome sequencing to obtain draft genomes of six lung isolates cultivated from BALF that represented the streptococcal diversity, using short read sequencing on the Illumina platform (Dataset S1, S2, S3). For species identification, we used both phylogenetic and biochemical approaches. Firstly, we performed genome-based identification using digital DNA-DNA Hybridization (dDDH)(19) on Type Strain Genome Server (TYGS) webtool(20). From this, we obtained species-level identification, the reference genomes of closest type strains and an outgroup taxon spanning a wide phylogenetic diversity (Table S1, S2, Dataset S4, S5). Additionally, we also obtained Genome BLAST Distance Phylogeny (GBDP)-based and full-length 16S rRNA-based phylogeny (Figure 1A, Figure S1B). Secondly, we performed pairwise whole genome Average Nucleotide Identity (FastANI(21)) including the reference genomes and outgroup taxon (Figure 1B, Figure S2A). Thirdly, for further precision, we generated single-copy core gene phylogeny (Figure 2A, Figure S2B). Fourthly, we identified lung isolates using routine clinical microbiology, which included optochin resistance test, Matrix-Assisted Laser Ionization Time-Of-Flight (MALDI-TOF)-based protein spectral analysis and hemolysis (Table S3, Figure S2B, C). Finally, we performed standardized biochemical and metabolic panel tests (Strep API20, Biomerieux) for phenotypic characterization (Table S4, Figure S3). All lung isolates were confirmed to be Viridans Group Streptococci (VGS, Figure S2B, C)(22). Phylogenetic analysis suggested that five out of six isolates represent potential new species. (i) *Streptococcus* isolate sp. nov. P2E5 is a novel species within the *S. mitis* group, occupying an intermediate phylogenetic position between the *S. mitis* (ANI 93.26%) and *S. pneumoniae* (ANI 92.72%) clades (Figure 1A, B, 2A). This isolate exhibits a typical α-hemolysis, proteomic analysis identified it as *S. mitis* whereas biochemical analysis indicated to *Gemella haemolysans.* (ii) *Streptococcus* isolate sp. nov. P2D11 is a new species within the *S. salivarius* group(23, 24). Interestingly, this isolate does not display any hemolysis (ɣ-hemolysis) and generated a unique biochemical pattern unlike a typical *Streptococcus* (Table S3, S4). (iii) *Streptococcus* isolate sp. nov. 369.3 is novel species that phylogenetically similar to *S. bovis* group and exhibits weak α-hemolysis (Figure 1, 2, S2, Table S3, S4). However, proteomic analysis indicated *S. parasanguinis* and *S. australis* (Table S3) and biochemical identification show similarities to *Gemella morbillorum* with low discriminatory power (Table S4). (iv, v) *Streptococcus* isolates sp. nov. P3B4 and P3D4 represent two closely related new species with phylogenetic and spectral similarities to *S. parasanguinis*, displaying typical α-hemolysis (P3B4 exhibited weak hemolysis). However, these two isolates exhibit biochemical and metabolic characteristics similar to *S. mitis group.* Lastly, *Streptococcus* P3E5 was identified as a new subspecies of *S. constellatus*, characterized by β-hemolysis, indicative of this specific this species. We henceforth, referred it as *S. constellatus* spp. nov. P3E5.

**Figure 1.**
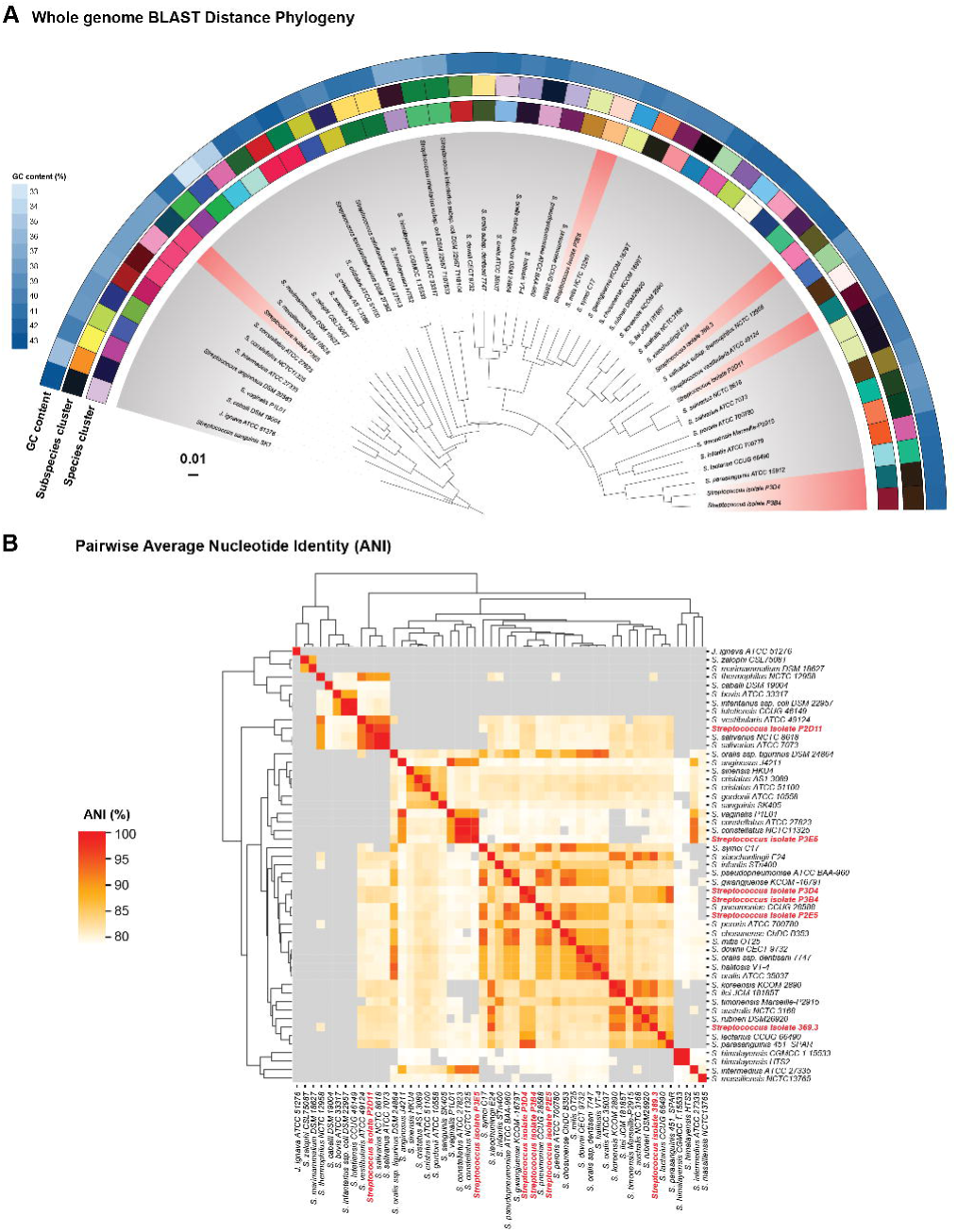
Whole genome phylogeny reveals novel streptococci from human distal lung microbiota. Comparison of human distal lung streptococci (red gradient) to closely related type strains (gray gradient) in the TYGS database. **A.** Whole Genome BLAST Distance Phylogeny (GBDP) using FASTME, where colored boxes represent species and subspecies clusters, and blue-colored gradient boxes represent GC content (%). **B.** Heatmap showing pairwise Average Nucleotide Identity (ANI %) between human distal lung streptococci (red gradient) to closely related type strains (gray gradient). ANI values > 80% were colored as grey in the heatmap.

**Figure 2.**
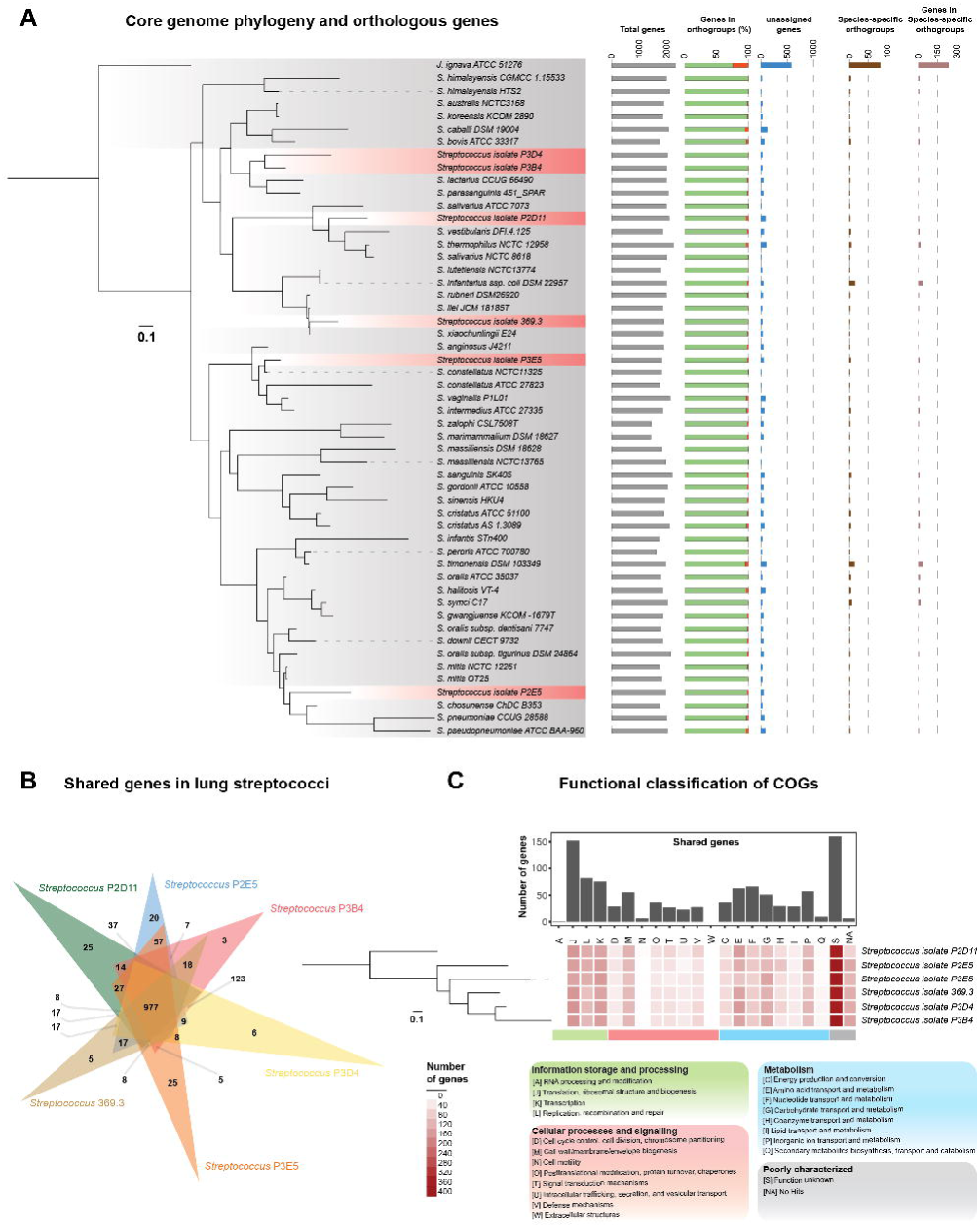
Core genome phylogeny and orthologous gene content of human distal lung streptococci A. Single copy core-genome phylogeny and orthologous gene analysis of lung streptococci (red gradient) and closely related type strains (gray gradient). Maximum-likelihood tree computed by FastTree on concatenated amino acid sequences of 315 single-copy core proteins using LG + CAT substitution model. Bar graphs show total number of genes (gray bars), percentage (%) genes in orthogroups (green and red stacked bars), number genes that weren’t assigned to orthogroups (blue bars), number of specific-specific orthogroups (dark brown) and number of genes in each (light brown). **B**. Venn diagram showing 977 shared proteins between lung streptococcal isolates generated using OrthoVenn2. **C**. Heatmap showing functional classification of proteins from lung streptococcal isolates using eggNOG depicted according to maximum-likelihood phylogeny computed by RAxML on concatenated amino acid sequences of 957 single-copy core proteins. Bar graphs show number of shared genes (y-axis) in each functional category (x-axis, colored strips).

### Pangenome analysis sheds light on gene content and evolutionary relationship amongst lung streptococci

To understand the gene content of human lung *Streptococci* when compared to the reference genomes, we combined total proteins from 47 reference bacteria (Dataset S6) and 6 isolates (Dataset S7) to perform orthologous gene analysis using OrthoFinder(25). Overall in 53 genomes, we found 5232 orthogroups accounted for 97.7% (99,280/101,643) of all proteins along with 150 strain-specific orthogroups (2.8%) (Figure 2A, Dataset S8). Core genes represented 493 orthogroups (9.42%) with 315 single-copy core genes, which we used to construct a maximum likelihood evolutionary tree (Figure 2A) that corroborated the whole genome-based phylogeny (Figure 1). Next, we investigated the gene content of the Streptococcal isolates in comparison to each other. We combined all proteins from 47 reference genomes to construct a custom *Streptococcus* pan-proteome (Pan-Strep) database (Dataset S9) and compared each isolate to this using OrthoVenn2(26)(Dataset S10). The Pan-Strep database contained 31726 orthologous (169,727 proteins) with majority of the proteins present in lung isolates with few exceptions. *Streptococcus* isolate sp. nov. P2E5, P2D11, P3B4, P3D4, 369.3 and *S. constellatus* spp. nov. P3E5 contained 1970, 2086, 1988, 2024, 1833 and 1891 genes belonging to 1780, 1846, 1872, 1883, 1757 and 1696 gene clusters respectively. Furthermore, these isolates shared 977 genes including 957 single-copy core genes (Figure 2B). Interestingly, all isolates contained only 1 unique gene cluster each with none found in *Streptococcus* isolate sp. nov. P2E5. For functional categorisation, we performed Clusters of Orthologous Groups (COG) analysis in individual isolates and shared genes using eggNOG mapper(27). We found 22 COGs including 20 with known functional groups and 2 with unknown function (Figure 2C, Table S5, Dataset S11). Majority of the genes belonged to the COG category of unknown function (S) followed by translation, ribosomal structure and biogenesis (J), transcription (K) and Replication, recombination and repair (L). The most abundant cellular process was cell envelope biogenesis (M), and the most abundant metabolic genes were responsible for amino acid metabolism (E), carbohydrate metabolism (G) and inorganic ion metabolism (P). Contrastingly, cell motility (N), RNA processing and modification (A), extracellular structure (W), intracellular trafficking, secretion and vesicular transport (U) and lipid transport and metabolism (I) were not prevalent. Although phylogenetically different, shared most of the COGs indicating similarities in basic cellular and metabolic functions.

### Metabolic and functional analysis of distal lung streptococci provide insights on the lung microbial ecosystem

Next, we predicted metabolic functions and macromolecular machineries using a custom rule-based based pipeline that included the GapMind, dbCAN and MacSysFinder tools (28–32) to comprehensively investigate the common catabolic and biosynthetic pathways, secretion systems, bacterial competence and carbohydrate-active enzymes (CAZymes)(33)(Figure 3, Table S6). The most prevalent mechanism for carbon catabolism in all streptococci including the lung isolates was the Embden-Meyerhof-Parnas (EMP) pathway. This was followed by Pentose Phosphate pathway (PPP) and Entner-Doudorrof (ED) pathway amongst majority of streptococci we tested. However, in majority of the bacteria, we observed either complete absence or incomplete canonical TCA cycle and oxidative phosphorylation (OXPHOS).

**Figure 3.**
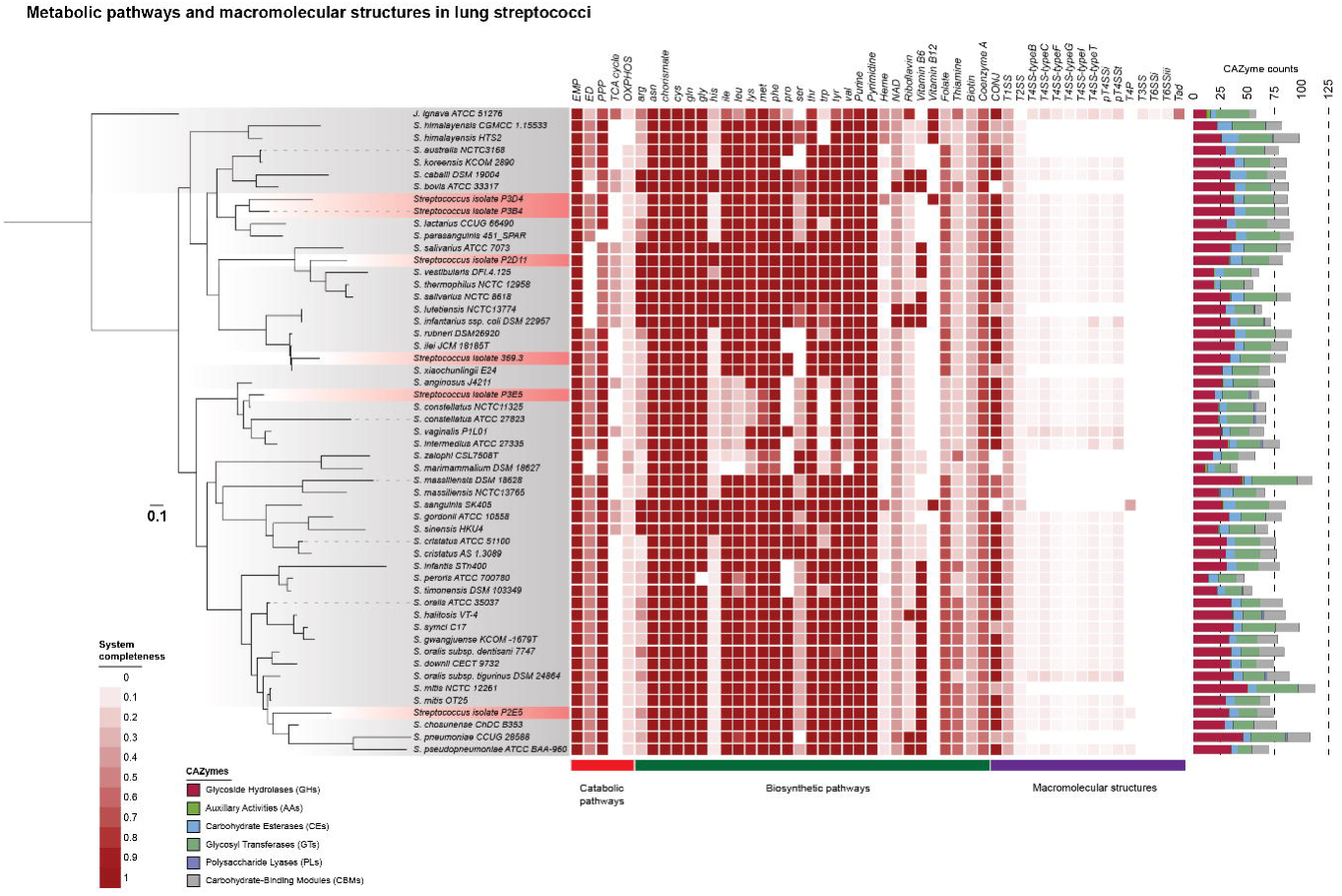
Function and metabolic prediction predictions of human distal lung streptococci. Heatmap showing completeness of predicted pathways depicted according to maximum-likelihood phylogeny computed by FastTree on concatenated amino acid sequences of 315 single-copy core proteins using LG + CAT substitution model in individual human distal lung streptococci (red gradient) and closely related type strains (gray gradient). A custom rule-based pipeline was used to predict metabolic pathways; catabolic (red stripe), biosynthetic pathways (green stripe) and macromolecular systems (purple stripe). Stacked bar charts show counts (y-axis) of Carbohydrate-active enzymes (CAZymes) in individual genomes (x-axis).

All lung isolates except *Streptococcus* isolate sp. nov. P2D11 possessed complete PPP, which was consistent within the *S. salivarius* group. We further investigated the number of carbohydrate-active enzymes (CAZymes, Figure 3, Figure S4A, Table S6, Dataset S12) and individual capacity to ferment sugars in culture (Figure S3). In total, the Pan-Strep database contained 5705 CAZymes subdivided into 6 families (Figure S4A). Compared to *S. pneumoniae* (108 CAZymes) that could ferment D-lactose, D-raffinose and D-trehalose. *Streptococcus* isolate sp. nov. P2E5 (mitis group, 74 CAZymes) and *Streptococcus* isolate sp. nov. 369.3 (*S. bovis* group, 85 CAZymes) exhibited no sugar fermentation despite having GH1, 4 family of Glycosyl Hydrolases (Figure S4B). *Streptococcus* isolate sp. nov. P3B4 (88 CAZymes) and P3D4 (87 CAZymes) have a similar biochemical and metabolic profiles with D-lactose and D-raffinose fermentation capability, with the latter additionally fermenting D-sorbitol. *Streptococcus* isolate sp. nov. P2D11 (83 CAZymes) ferments D-mannose, D-sorbitol and D-lactose. *S. constellatus* novel. spp. P3E5 (61 CAZymes) was only able to ferment D-trehalose.

Next, we show that all isolates possessed pathways for the biosynthesis of most amino acids with a few exceptions (Figure 3, Table S6). All bacteria possess chorismate biosynthesis pathway (*aro*G, *aro*B, *aro*D, *aro*E, *aro*L, *aro*A, *aro*C), which can serve as an intermediate for biosynthesis of essential amino acids. In addition, all streptococci had complete pathways for nucleotide biosynthesis (purines and pyrimidines). However, most streptococci including lung isolates lacked the genes necessary for biosynthesis of electron acceptors and mediators such as Heme, NAD^+^, Coenzyme A and vitamin biosynthesis with interesting exceptions. Unlike *Streptococcus* isolate sp. nov. P3B4, *Streptococcus* isolate sp. nov. P3D4 can synthesize Vitamin B12. In addition, *Streptococcus* isolate sp. nov. P2D11 possessed the capability for Vitamin B6 biosynthesis consistent with the *S. salivarius* group.

We also investigated the presence of macromolecular structures such as secretion systems, diversity of competence and DNA uptake complexes in human lung commensal Streptococci. As expected, the most prevalent of these multi-protein complexes found in streptococci were the competence (com) proteins (Figure S5) responsible for natural competence, DNA uptake and transformation. These included competence stimulation peptides (CSP) and export protein (ComB, ComC)(34, 35) and the major response regulator ComX(36). Two types of DNA uptake complexes were found across all genomes: ComE proteins (comEA, EB, EC)(37) and the ComF proteins (38) and the ComG pilus-like proteins (comGA, GB, GC, GD, GE, GF, GG)(39, 40).

Although no complete secretion systems were presentin all genomes (Figure 3, Figure S5, Table S6), we still found Type IV secretion system (T4SS) proteins involved in conjugation(41). More specifically, proteins that we considered mandatory for a functional conjugative process were the ATPase complex system (VirB4)(42), coupling proteins (T4CP1, T4CP2)(41) and the type 4 toxin co-regulated pilus (TCP) subunit system (TcpA)(43) were present in all lung isolates. The accessory system including relaxases (MOBs)(44) were more variable across genomes with 5 MOB families (MOB_B_, MOB_C_, MOB_Q_, MOB_T_, MOB_V_) detected (Figure S5). Hence, we classified these as conjugation system proteins (CONJ)(28).

### Occurrence and prevalence of antimicrobial resistance and virulence factors in human distal lung streptococci

Here, we investigated the presence of antimicrobial resistance (AMR) genes and virulence factors in the lung *Streptococcus* isolates using the ABRicate tool(45) (Figure 4A, Table S7, S8). We also corroborated this with antibiotic susceptibility assays for all six isolates following EUCAST protocols that includes both disk diffusion assays and MIC tests (Table S2).

**Figure 4.**
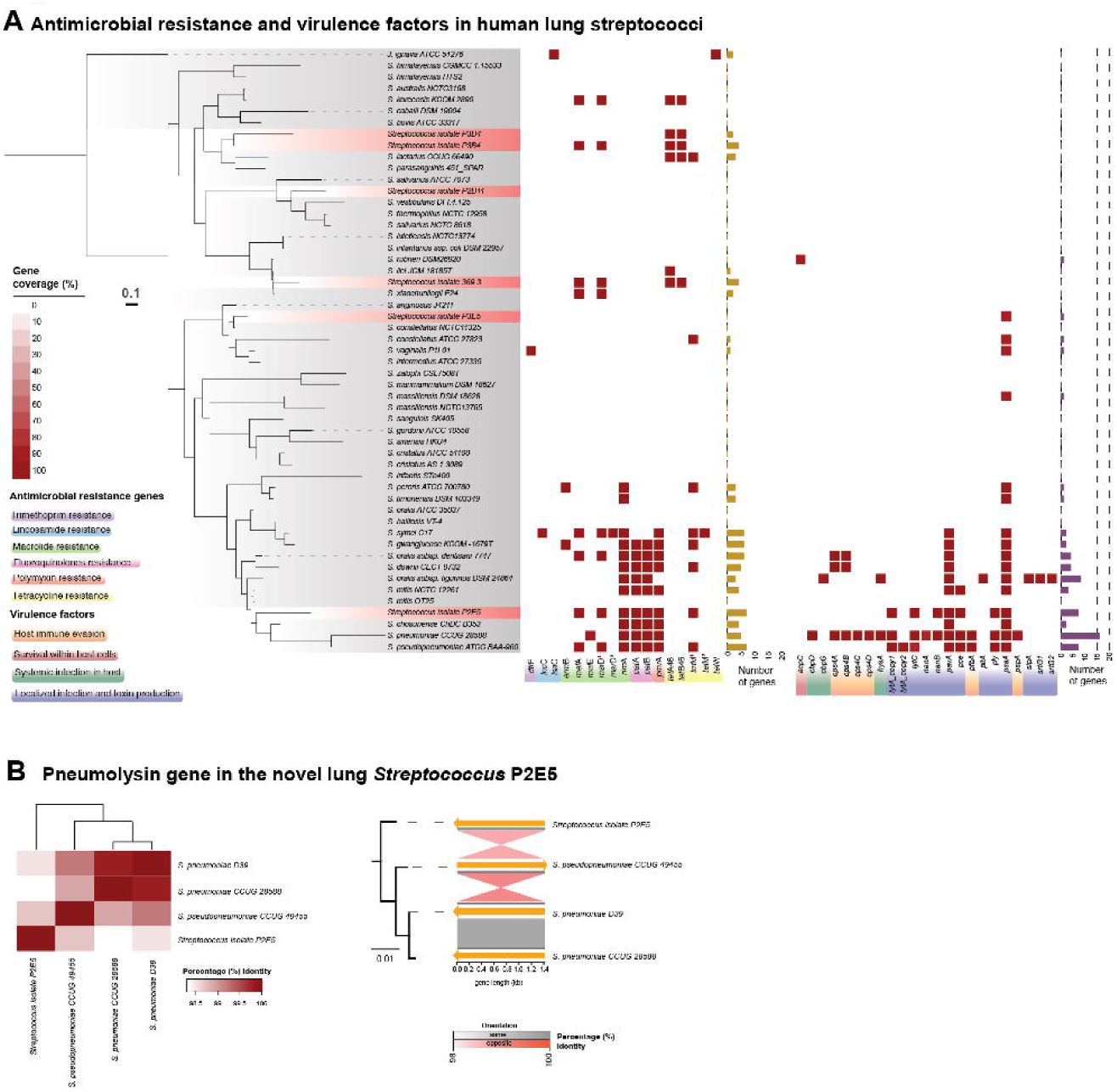
Prevalence of antimicrobial resistance and virulence factors in human distal lung streptococci. A. Heatmaps depicting coverage of antimicrobial resistance (left panel) and virulence gene (right panel) revealed by ABRIcate depicted according to maximum-likelihood phylogeny computed by FastTree on concatenated amino acid sequences of 315 single-copy core proteins using LG + CAT substitution model in individual human distal lung streptococci (red gradient) and closely related type strains (gray gradient). Bar graphs show number of antimicrobial resistance (yellow) and virulence (purple) genes (y-axis) in each category (x-axis, colored strips). **B.** Heatmap depicting pairwise distance matrix after alignment generated using MUSCLE, showing percentage identities of Pneumolysin protein between *Streptococcus* isolate P2E5 and pneumococci. Maximum-likelihood phylogeny of Ply proteins computed by by RAxML and gene plot for genomic coordinates and synteny of *ply* genes generated by pyGenomeViz. This compared the percentage identity (shading depth) and relative gene orientation (grey for links with the same direction, pink for reverse direction). This compared the percentage identity (shading depth) and relative gene orientation (grey for links with the same direction, pink for reverse direction).

Comparison with the MEGARes database(46) revealed AMR genes in 27 reference genomes and 4 isolates (*Streptococcus* isolates isolate sp. nov. 369.3, P3D4, P3B4 and P2E5), including multiple variants and /or copy numbers conferring resistance to 6 classes of antimicrobials (Figure 4A). The most prevalent AMR was against Macrolides (66.7%; 18/27 genomes) followed by Tetracyclines (63%; 17/27) and Fluoroquinolones (37%; 10/27 genomes). Macrolide resistance genes (*mef*A and a single copy of *msr*D)(47, 48) were observed in 3 isolates apart from *Streptococcus* isolate sp. nov. P3D4. This was confirmed by antibiotic susceptibility assays (Table S2) where *Streptococcus* isolate sp. nov. P3B4 was resistant to Erythromycin (11 mm, MIC breakpoint = 4 mg/L), whereas *Streptococcus* isolate sp. nov. P3D4 was sensitive (27 mm). Although we didn’t find any Lincosamide resistance genes (*lnc*C, *lsa*C) in lung isolates, we tested susceptibility towards Clindamycin and performed D-test to distinguish between M- and MLS_B_-phenotype of macrolide resistance(48, 49). All isolates were sensitive to Clindamycin and showed no MLS_B_-phenotype, indicating only the presence of M-phenotype in human distal lung streptococci.

Tetracycline resistance genes were found in 4 isolates with *tet*A46 and *tet*B46(50) present in 3 isolates i.e., *Streptococcus* isolates sp. nov. 369.3, P3B4 and P3D4 and only one copy of *tet*M in one isolate i.e., *Streptococcus* isolate sp. nov. P2E5. Interestingly, all four isolates showed marginal resistance in diffusion assays and none in MIC tests. In addition, *Streptococcus* isolate sp. nov. P2D11 and P3E5 neither harboured the genes nor exhibited resistance phenotype. Interestingly, although we didn’t find genes related to beta-lactam resistance, all lung isolates exhibited Oxacillin resistance (disk diffusion) and 3/5 isolates were resistant to Benzylpenicillin (MIC breakpoint = 0.25 mg/L, Table S2).

Virulence factor analysis using the VFDB(51) database revealed 20 genomes (2 isolates and 18 references) harboring diverse virulence-related genes (Figure 4A). The highest number were observed in *S. pneumoniae* with *psa*A encoding for pneumococcal surface adhesin A, which plays a role in general and localized infection with *Streptococcus*(52, 53) the most prevalent across genomes. In one of the most interesting findings, we found pneumolysin (*ply*) gene, autolysin-encoding gene (*lyt*A) and pneumococcal surface adhesin A (*psa*A) in the novel isolate *Streptococcus* P2E5, the closest match to the most prevalent *Streptococcus* in human lung(18) and *psa*A in *Streptococcus* isolate sp. nov. P3E5. These genes are generally used to identify *S. pneumoniae*(54). Upon phylogenetic analysis, we show that *Streptococcus* isolate sp. nov. P2E5 Ply protein is similar to that of *S. pneumoniae* CCUG 28588 (98.3% identity) and *S. pseudopneumoniae* CCUG 49455 (98.7% identity) and type strain *S. pneumoniae* D39V (98.5% identity) (Figure 4B, Dataset S13). These results indicate the general abundance of AMRs but scarcity of virulence genes in VGS, including the lung isolates. Therefore, *Streptococcus* isolate sp. Nov. P2E5 is not only phylogenetically intermediate to *S. pneumoniae* and *S. mitis* but also in terms of virulence factors.

### Lung streptococci exhibit variable capsular diversity

Capsular polysaccharides common in commensal streptococci(55). However, we didn’t observe all capsule genes in our virulence factor analysis. Hence, we investigated the presence of capsule genes in all genomes by protein BLAST against our Pan-Strep database using the prototypical *S. pneumoniae* D39 (Sp D39, serotype 2) capsule (*cps*) locus i.e., the 17 genes located in between the *dex*B and *ali*A genes, as the query sequence followed by phylogenetic analysis of the matching genes (Figure 5A, Dataset S14). In addition, we also performed serotyping of the lung isolates by Quellung’s test(56). Out of 53 genomes, 36 contained one or more capsular genes (Figure 5B, C) with the lowest (2 proteins) found in *Streptococcus pseudopneumoniae* ATCC BAA-960 and highest (22 proteins) in *Streptococcus salivarius* NCTC 8618. All six isolates possessed capsular proteins, which were a subset of Sp D39 (17 proteins) *cps* genes; P2D11 (12 proteins), P2E5 (13 proteins), P3E5 (11 proteins), P3B4 (12 proteins), P3D4 (11 proteins) and 369.3 (13 proteins).

**Figure 5.**
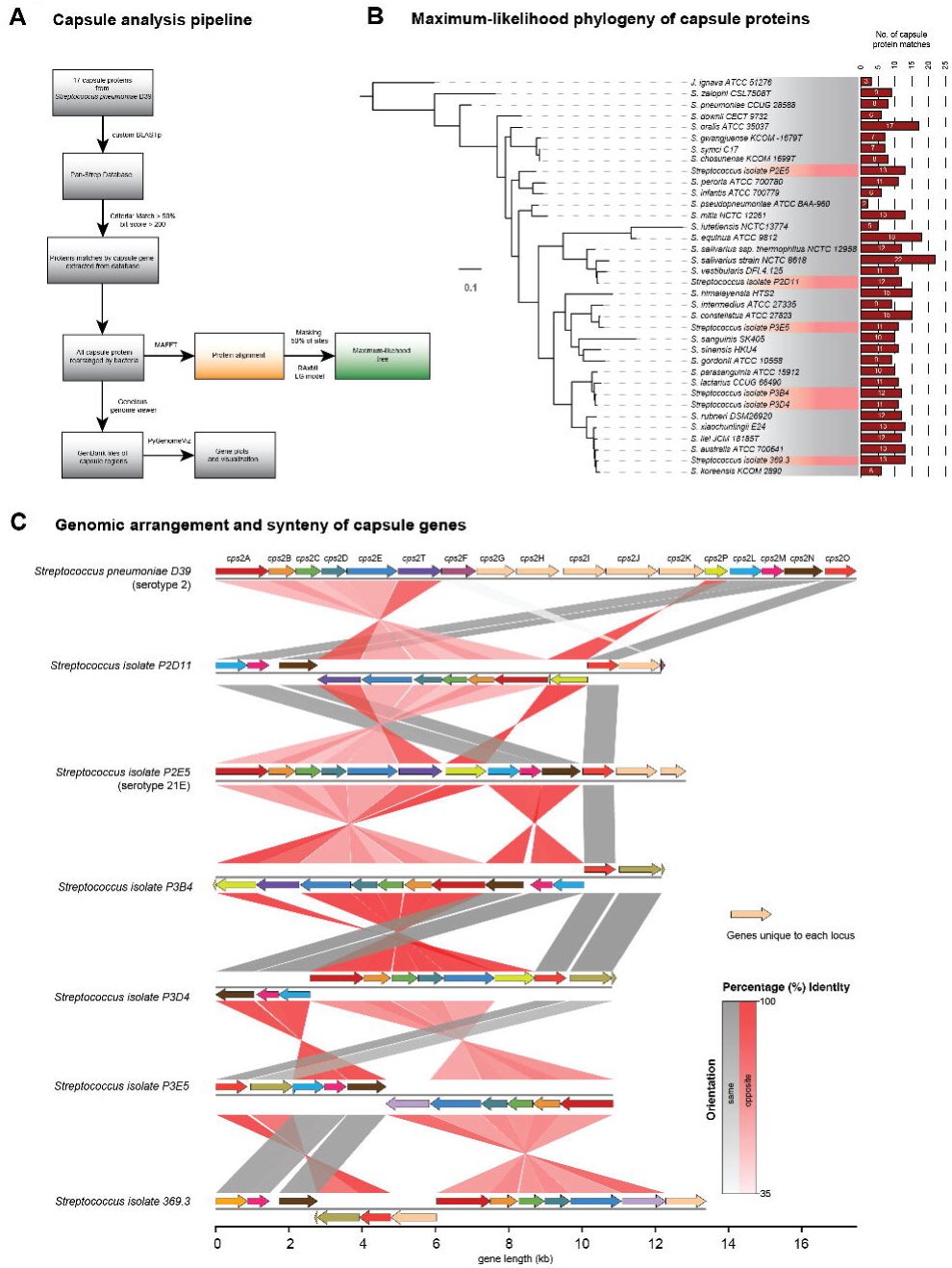
Investigation of Streptococcal capsular polysaccharide synthesis and its relative genomic arrangement. A. Workflow showing our capsular analysis pipeline from BLAST search to phylogeny. **B.** Maximum-likehood phylogeny computed computed by by RAxML after alignment of capsular proteins found in individual human distal lung streptococci (red gradient) and closely related type strains (gray gradient). Bar graphs show number of capsule protein BLAST hits (y-axis) in individual genomes (x-axis). **C.** Comparison of capsular polysaccharide synthesis (*cps*) genes (colored arrows) in lung isolates to the prototypical *S. pneumoniae* D39 capsule operon. Gene plot generated by pyGenomeViz for comparing genomic coordinates and synteny with percentage identity (shading depth) and relative gene orientation (grey for links with the same direction, pink for reverse direction).

The Sp D39 genes absent in isolates encoded for GTB-type glycotransferase superfamily of proteins(57) (Cps2G and CpsI), Capsular synthesis protein(58) Cps2H (*cps*2H), MATE-family protein(59) Cps2J (*cps*2J) and UDP-glucose 6-dehydrogenase Cps2K(60) (*cps*2K). However, unique capsular genes were also found in *Streptococcus* isolate sp. nov. P2D11 (hypothetical protein glycotransferase 1 family, protein ID EKHPBGBN_01095), *Streptococcus* isolate sp. nov. 369.3 (diaminopimelate decarboxylase; *lys*A, UDP-galactopyranose mutase; *glf*) and *Streptococcus* isolate sp. nov. P2E5 (UDP-galactopyranose mutase; *glf*2, UTP-glucose-1 phosphate uridylyltransferase; *cug*P). Interestingly, only *Streptococcus* isolate sp. nov. P2E5 tested positive for Quellung’s test and was characterized to be serotype 21E. Hence, we compared its capsule genes with that of *S. pneumoniae* 546/62 (Sp 546/62, reference for serotype 21) along with Sp D39 (serotype 2). We show that 11/13 capsule genes in P2E5 were high similarity to both Sp 546/62 and Sp D39 (Figure S6), indicating similarity to both serotypes.

### Core genome phylogeny reveals evolutionary relationship between lung, oral and type strains of Streptococci

Although being of oral and supraglottic origin, the distal lung microbiota distinct (9– 11). However, there is no study showing comparison at whole genome level. This prompted us to investigate the genetic proximity of the Streptococcal isolates cultivated from BALF and reference genomes from TYGS to that of the human oral microbiome. We used full-length 16S rRNA genes from the isolates and performed BLASTN against all genomes in the expanded Human Oral Microbiome Database (eHOMD)(61). This resulted in 47 representative genomes (best hits; > 97%16S rRNA gene identity), which we combined with lung streptococci and TYGS reference genomes to perform core genome phylogeny (Figure 6, Table S9, Dataset S15). Overall, we did not observe body-site dependent pattern emerging rather all Streptococci were phylogenetically distributed regardless of the origin. Remarkably, lung streptococci stood out as phylogenetically distinct with one exception (Figure 6). *Streptococcus* isolate sp. nov. P2E5 was phylogenetically distinct with no closely related bacteria in the oral repertoire (Figure S7A). *Streptococcus* isolate sp. nov. P2D11 was closely related and intermediate to oral *S. salivarius* and *S. vestibularis* genomes in HOMD (Figure S7B). Interestingly, *S. constellatus* spp. nov. P3E5 was phylogenetically closer to oral *S. intermedius* but still within the *Streptococcus anginosus* group (Figure S7C). The phylogenetic placement of *Streptococcus* isolate sp. nov. 369.3 didn’t change in relation to oral streptococci (Figure S7C). Finally, *Streptococcus* isolate sp. nov. P3B4 and P3D4 were observed to be phylogenetically distinct from both reference genomes and oral streptococci (Figure S7D). These results strengthen our findings of novel Streptococci and supporting the claims that lung microbiota is phylogenetically distinct from oral microbiota.

**Figure 6.**
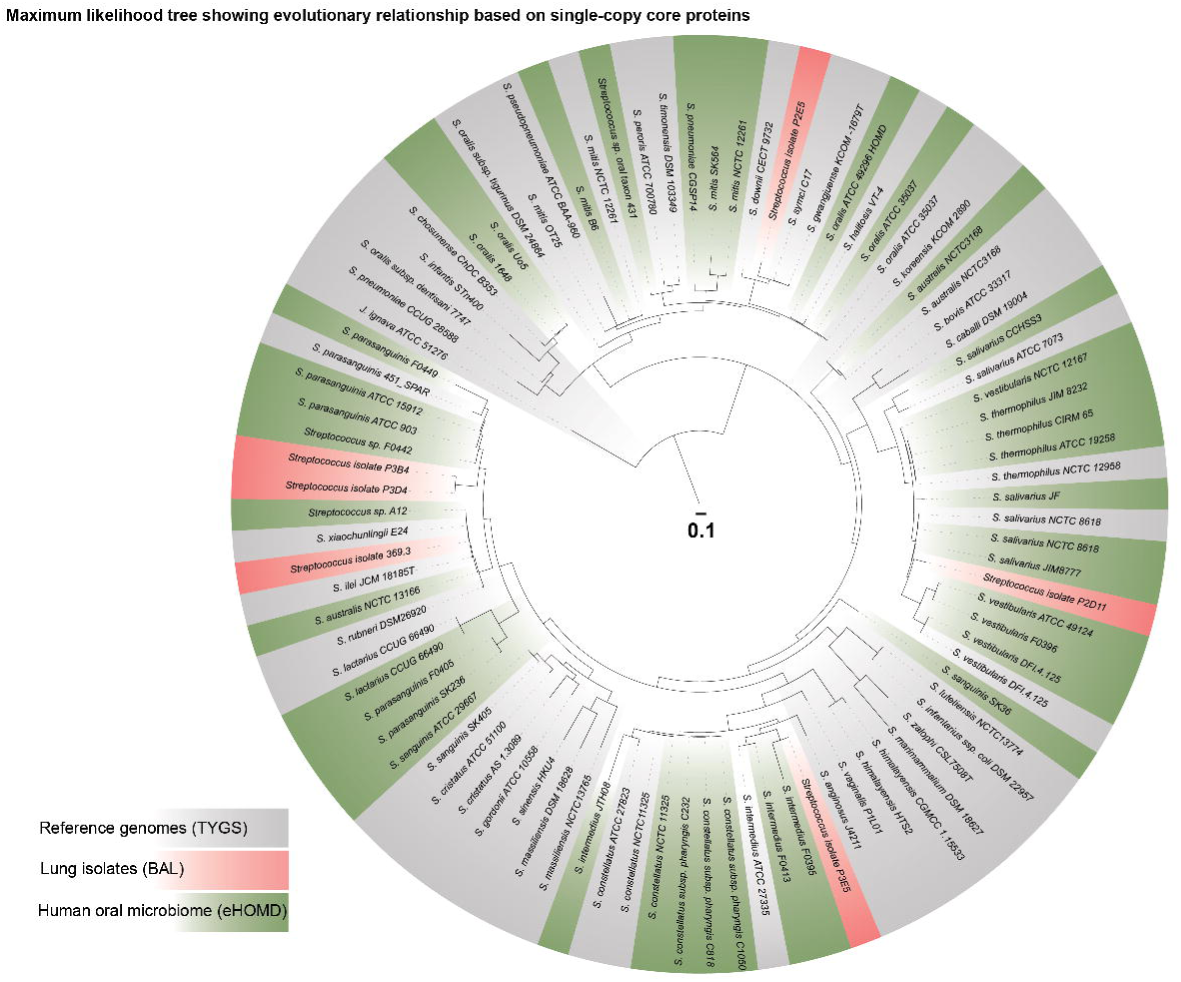
Phylogenetic comparison of human oral and distal lung streptococci. Single copy core-genome phylogeny comparing lung streptococcal isolates cultivated from BAL (red gradient), Reference genomes from TYGS (gray gradient) and human oral streptococci from eHOMD (green gradient). Maximum-likelihood tree computed by FastTree on concatenated amino acid sequences of 26 single-copy core proteins using LG + CAT substitution model.

## Discussion

Microbiota of the healthy lung is primarily derived from the oral and supraglottic niche(5, 8, 9, 12, 62, 63). This is also reflected in the lung post-transplant, where the oral taxa-dominant microbiota profile was associated with normal lung function and homeostasis(18, 64). Amongst all, Streptococci are the most spatiotemporally ubiquitous in oropharyngeal niche, upper and lower respiratory tract in healthy lung and allografts (5, 9, 11, 62–66).

Likewise, in our previous study we have established an important resource called LuMiCol containing several lung bacterial isolates that match top lung taxa revealed in amplicon sequencing(18). We also showed that *Streptococcus* is the most phylogenetically diverse and abundant genus in distal lung microbiota. However, due to limited resolution from amplicon sequencing, deeper genetic diversity in terms of specific species or strains were not known. Here, using robust phylogenomic analysis, comparative genomics and *Streptococcus*-specific phenotyping, we characterized 6 different novel streptococcal isolates (Figure 1, 2, S3), which belonged to the highly heterogenous VGS and are evolutionary intermediates to already existing human-associated Streptococci, including both commensal and pathogenic species. We also categorized these into species groups whenever possible, which can be inconsistent (67). For example, *Streptococcus* isolate sp. nov. 369.3 is genetically related to *S. bovis* group II/1 (mannitol negative and beta-glucuronidase negative, Figure1, 2, Table S4) but shows biochemical similarity to the Nutritionally Variant Streptococcus (NVS) *G. morbillorum* (68). In addition, *Streptococcus* isolate sp. nov. P2E5 has both phenotypic features of the *S. mitis* group (Table S3) and biochemical features of *G. haemolysans* (Table S4). These observations along with the fact these novel isolates possess known orthologs upon comparison to the Pan-Strep database indicates intra-genera rather than an inter-genera gene transfer. Despite being high-quality these are not closed genomes and information on complex genetic structures might be missing especially considering the high genetic variation in *Streptococcus*. Hence, a combinatorial approach using short- and long-read sequencing should be the next appropriate step.

Human lung microbiota is primarily composed of facultative or obligate anaerobes, including the streptococci reported in this study (18). However, little is known about the microbial metabolism in the deep lung. Streptococci not only represent a larger subset of resident bacteria but are also temporally and spatially the most prevalent genus(11, 18). Hence, its metabolic capabilities can provide crucial information on common catabolic and biosynthetic pathways within the lung microbiota. Although several genes for utilization and transport of sugars were present, there was a lack of canonical TCA cycle (Figure 3, Table S5). Additionally, the presence acetyl-CoA - Pyruvate/Lactate interconversion pathway (*ack*A, *pta* and *ldh*) (69) indicate a preference for anaerobic metabolism, which is in line with low glucose availability in airway epithelia(70). Hence, there might be two plausible pathways: the acetate-driven alternative TCA cycle (71) or pyruvate fermentation(69, 72).

The presence of complete pathways for acetate metabolism indicates its central role in the lung environment, which may be contributed mainly by commensal Streptococci (69, 73). Additionally, it is also an important short-chain fatty acid with immunomodulatory function in host gut and lung(74, 75) and shown to enhance killing of major lung pathogen *S. pneumoniae* by macrophages(76). However, these were mostly predictions, and we still lack information on nutritional preferences, which should be shown large-scale growth analysis on individual carbon sources.

Macromolecular structures in bacteria perform important functions in interacting with its environment. We revealed the presence multi-protein complex systems involved in bacterial competence, extracellular DNA uptake. All lung isolates harbor complete pathways for natural competence i.e., Com proteins including the pheromone peptides and regulators responsible for natural competence and extracellular DNA uptake complexes: ComE, F and G proteins (Figure S5)(34–40). However, this should be supported by further experimental induction of competence followed by DNA uptake(77). Lung streptococci also possess conjugative abilities shown by the presence of the VirB4)(42), TcpA)(43), T4CP1 and T4CP2(41) and may exchange genetic material with other genera in the community acquiring new traits.

Antibiotic resistance and virulence factors were mostly found in the mitis group VGS and pneumococci (Figure 4). Previously studies showed that antibiotic resistance is widespread in VGS and other human associated streptococci(78). Amongst the lung isolates, *Streptococcus* isolate sp. nov. P2E5 had most number with 7 genes (Table S6) and resistance pattern similar to other members of *S. mitis* group. As previously described for tetracycline resistance in oral streptococci(50), *te*tM encoding for ribosome protecting proteins was more common in our lung isolates than *tet*AB encoding for efflux pumps. Remarkably, presence of these genes did not manifest into phenotype (Table S6) apart from *Streptococcus* isolate sp. nov. 369.3. This could be due the requirement of additional genes or a result of altered gene regulation. Hence, it is challenging to conclude due the limitation that Tetracycline resistance in VGS is ill-defined by EUCAST due to insufficient evidence. (75). Interestingly, majority of lung isolates were resistant to the narrow spectrum beta-lactam Oxacillin, although we couldn’t report resistance genes. However, remains inconclusive without investigating penicillin binding proteins (PBPs), which are crucial in conferring beta-lactam resistance to streptococci(79). Also, pneumococcus-specific virulence factors such as *ply*, *lyt*A and *psa*A was found in *Streptococcus* isolate sp. nov. P2E5, which tested positive of pneumococcal polysaccharide capsule (serotype 21E) (Figure 5). However, it had less genes when compared to known serotype 21 reference genome Sp 546/62 (Figure S6). The unique proteins in P2E5 such as UDP-galactopyranose mutase (*glf*2) and UTP-glucose-1 phosphate uridylyltransferase (*cug*P) may contribute to the specific capsule biosynthesis. However, this requires further investigation using other tests like immunodiffusion test(80) and heterologous expression of these genes to confirm its contribution. Remarkably, this isolate matched to the most prevalent *Streptococcus* in the human lung, which is associated with good lung function and immunological balance. Presence of pneumococcal capsule serotypes has been reported in VGS and other human associated streptococci(60, 81). But P2E5 is unique as it an evolutionary intermediate with clear features of pneumococcus and *S. mitis* (Figure 1A, B, 2A). This phenotypic diversity and intermediary features amongst lung streptococci along with the presence of functional machineries for horizontal gene transfer may indicate a high degree of genetic exchange in the lung microbial environment. Previous studies have shown despite finding its origin in oral niche, the structure and composition of lung microbiota is distinct. Here, we provide genome-level evidence for the first time and show that lung microbiota remains phylogenetically distinct when compared to oral isolates from eHOMD.

Hence, to our knowledge, this is the first study to genome sequence novel lung bacterial isolates and perform comparative genomics to reveal crucial genetic, metabolic and evolutionary information filling the knowledge gaps in the field of microbial ecology of the human distal lung.

## Materials and Methods

### Sample collection and ethics

Sampling was performed and anonymized as previously described(18). The sampling via bronchoscopy was performed on individuals post lung transplant. Bronchoalveolar lavage fluid (BALF) was cultivated at random on different media and at different oxygen conditions. This sampling was approved by the local ethics committee (“Commission cantonale (VD’ d’éthique de la recherche su’ l’être humain – CER-VD”, protocol number 2018-01818) with written informed consent.

### Data and code availability

All sequencing raw data were submitted to NCBI Short Read Archive under the BioProject PRJNA1001255. Individual isolates were submitted under different BioSamples i.e., SAMN36797456, SAMN36797455, SAMN36797454, SAMN36797453, SAMN36797452, SAMN36797451. Processed data and supplementary datasets were uploaded on zenodo under DOI 10.5281/zenodo.10220079. Processed data including metaQUAST files, FASTA sequences and annotation files. All codes and pipelines are available on the GitHub https://github.com/slipa17/Whole-genome-sequencing-and-comparative-genomics-of-human-lung-streptococcal-isolates. Details on any scripts (.sh files), workflows (.md /.Rmd or .R files) and parameters (.txt files), which are mentioned throughout can be be found on the GitHub page.

### Bacterial growth and media

For routine cultivation, all Streptococci were grown for 24 – 48 hours on Columbia agar (Oxoid, UK) with 5% defibrinated sheep blood (Thermo Scientific, USA) at 37°C in presence of 5% CO_2_ and 95 % relative humidity or in a vinyl anaerobic chamber with < 5 ppm O_2_ (Coy labs, USA) at 35°C with moisture control. For broth cultures, one or two isolated colonies were picked and inoculated in a polypropylene culture tube (with cap) containing 2 ml of Todd-Hewitt Broth (Oxoid, UK) supplemented with 0.5% Yeast Extract (Oxoid, UK). The cultures were incubated under the same conditions as mentioned above and strictly without agitation.

### Bacterial DNA isolation and genome sequencing

Some streptococcal isolates exhibited unusual physical properties upon growth on semi-solid media, which included dry, flaky colony texture, difficulty in resuspension in buffer and recalcitrance. Hence, a custom bacterial DNA isolation protocol was used. This process involved sequential lysis of bacteria using both enzymatic action and mechanical shearing followed by extraction using QIAamp DNA Mini Kit (QIAGEN, Germany). Bacteria were grown as broth cultures and harvested followed by resuspension in 200 μl of Gram-positive lysis buffer (20 mM Tris-HCl, pH 8.0, 2 mM EDTA, 1.2% Triton X-100) containing 1 mg/ml lysozyme and 100 μg/ml RNase A. The mixture was incubated for 30 min at 37°C with gentle agitation. After this, the volume was brought up to 500 μl with Gram-positive lysis buffer and the mixture was transferred to screw cap tubes in tubes containing 200 mg of 0.1-mm acid-washed zirconia beads and homogenized using a FastPrep-25 5G instrument (2 rounds of 30 s with the power set to 6), as previously described(28). This was followed by centrifugation at maximum speed for 5 minutes at room temperature. The debris-free supernatant was carried over to the QIAamp kit protocol, which involves incubation with Proteinase K followed by column-based extraction steps. Genomic libraries for Illumina sequencing libraries were prepared in-house using the Vazyme TruePrep DNA library preparation kit following manufacturer’s instructions. Multiplexing was performed using Nextera i7 adaptors. Sequencing was performed on the Illumina HiSeq 2500 instrument at the Genomics Technology Facility, University of Lausanne, Switzerland using two simultaneous lanes for avoiding lane-bias generating 150 bp pair-end reads.

### Bacterial genome assembly and annotation

Read quality control and trimming was performed using FastQC v0.11.9(82) and Trimmomatic v0.39(83) (parameters: PE-phred33 AllIllumina-Peadapters.fa:3:25:7 LEADING:9 TRAILING:9 SLIDINGWINDOW:4:15 MINLEN:60). SPAdes(84) (–careful option, v3.15.2) was used for *de novo* assembly of bacterial genomes using *run_spades.sh* (Dataset S2). Different parameters were used assess the quality of assemblies (spades_param.txt). Assemblies were evaluated for its quality and completeness using metaQUAST (Quality Assessment Tool for Genome Assemblies) v5.0.2(85) and checkM v1.0.13(86). All genome statistics and metaQUAST HTML report files were created for summaries of each assembly task (Dataset S1). Draft genome scaffolds were annotated using prokka v1.13(87) using *prokka.sh* (Dataset S3).

### Genome-based bacterial identification using Type Strain Genome Server (TYGS)

For identification of closely related *Streptococcus* species and Genome-based Distance Phylogeny (GBDP), the DNA FASTA files of the isolates were submitted to [TYGS](https://tygs.dsmz.de/)(20) web portal. This tool uses for genome and 16S rRNA BLAST with clusters species and subspecies to identify species and report nearest neighbours. The output includes genome and 16S rRNA based phylogenetic trees, which can be exported. Visualization of these trees and associated metadata was done using [iTOL](https://itol.embl.de/)(88) v6.8.1 webtool.

### Data extraction from public databases

All reference genomes from TYGS analysis (Dataset S4) including GenBank data files (.gbk, .gbff), annotation features (.gff), Nucleotide (.fna, .fa) and Protein (.faa) FASTA files were downloaded from NCBI FTP server using NCBI Datasets command-line tools (CLI), using *NCBI datasets and BLAST.md*. In case any genome assembly did not accompany translations or proper annotations, which were then annotated using PROKKA. Human oral isolates genomes were downloaded from [eHOMD Genomes](https://ehomd.org/genome/genome_table). For consistency and reproducibility of further analyses, all genomes were reannotated using prokka v1.13(87) using *prokka.sh* (Dataset S5, S14).

### Pairwise average nucleotide identity (ANI)

Pairwise Average Nucleotide Identity (ANI) including visualization between *Streptococcus* genomes was performed using *FastANI.md* that combined ANIcluster and FastANI command line tools (89, 90). The output files contained query genome, reference genome, ANI values, count of bidirectional fragment mappings, and total query fragments.

### Pangenome analysis and orthologous group

This involved three steps: 1. Creation of a Pan-Strep database: all proteins FASTA files available from reference genomes provided by the TYGS analysis(20) were concatenated to form the pan-proteome database, which was used both for orthology and BLAST database (Dataset S9). 2. Comparison of isolates to Pan-Strep database: OrthoVenn2 webtool(26) was used classify orthologous gene clusters in each isolate using Pan-Strep database as reference. The E-value and inflation value were set at 1 x 10^-5^ and 1.5 respectively. The distribution of the shared (only between isolates) and unique orthologous clusters, total protein count was represented as a Venn diagrams (Dataset S10). 3. Comparison of all genomes to each other: All protein FASTA files from genomes were analyzed together using Orthofinder v2.5.5(25) to obtain the single copy core orthogroups. This provides multiple statistics on the genetic contents including orthologs, xenologs and shared genes and single-copy shared genes (Dataset S8).

### Single copy core genome phylogeny

A list of single copy core (shared) genes (Dataset S7) used to extract corresponding protein FASTA from each genome and create individual FASTA files containing core genes using *Append_concatenate_extract_grep_Linux_log.md* and *Extraction_of_headers_fasta_sequences_and_splitted_files.md*. All proteins in each FASTA file were self-concatenated to create a single sequence FASTA with one header. All resulting sequences were aligned using MAFFT v7.52 command line tool(91) and Maximum-likelihood (ML) tree was computed using the FastTree v2.1.11(92) or RAxML v 8.2.12 command line tool (93). Visualization of these trees was done using [iTOL](https://itol.embl.de/)(88) v6.8.1 webtool.

### Multiple sequence alignment and phylogeny

All alignments (Protein or DNA FASTA sequences) were performed using MAFFT v7.52(91) command line tool and phylogenetic trees were constructed using FastTree v2.1.11 or RAxML v8.2.12 command line tool (93), unless and otherwise specified. The workflow *Alignment and phylogeny.md* was used that includes MAFTT for MacOSX (version 7.505) with the argumen–*‘--auto*’ for automatic detection of parameters. The output FASTA file containing the aligned sequences was masked (50% sites stripped) using Geneious Prime software (New Zealand) and exported as PHYLIP format. Maximum-likelihood (ML) trees was computed with the alignment file using either RAxML v8.2.12 (model ‘*PROTGAMMAAUTO*’, 100 rapid bootstrapping) or FastTree v2.1.11 (LG + CAT substitution model for protein or default -nt model for nucleotides). Visualization of these trees was done using [iTOL](https://itol.embl.de/)(88) v6.8.1 webtool.

### Comparison with genomes from expanded Human Oral Microbiome Database (eHOMD)

For this analysis, full-length 16S rRNA gene sequences of the isolates were used as queries to search on the [HOMD RefSeq BLAST Server](https://ehomd.org/genome/blast_sserver?type=refseq) (perform BLASTN v2.12.0, HOMD_16S_rRNA_RefSeq_V15.23.p9 database). The hits (>97% identity and >99% coverage) were selected and its genomes were downloaded from eHOMD. These genomes were reannotated using prokka v1.13(87) using *prokka.sh.* Comparison with the lung isolates and reference type strains from TYGS was performed by single-copy core genome phylogeny (Dataset S14). The protein FASTA files all sources were used to run Orthofinder v2.5.5 to obtain the single copy core orthogroups. These orthogroup FASTA files were extracted using *Append_concatenate_extract_grep_Linux_log.md* and *Extraction_of_headers_fasta_sequences_and_splitted_files.md* that rearranges these into single-copy core proteins according to each genome. These protein files were then aligned with MAFFT v7.52 and tree was constructed using FastTree v2.1.11 (LG + CAT substitution protein model). Tree visualization was done using [iTOL](https://itol.embl.de/)(88) v6.8.1 webtool.

### Clinical microbiology, identification and phenotyping

Species identification was done as described(94). In brief, routine bacterial identification was performed using a combination of MALDI-TOF, serotyping and functional assays at the Institute for Infectious Diseases (IFIK), University of Bern, Switzerland. Streptococci were grown on Columbia agar with 5% defibrinated sheep blood (Biomérieux) (CSBA plates) for 24 hours at 37°C in presence of 5% CO_2_. Single colonies were picked for MALDI-TOF analysis. In addition, bacterial were spread on CSBA plates as lawn cultures and Optochin disks (Sigma) was placed with a sterile forceps. The plates were incubated for 24 hours at 37°C in presence of 5% CO_2_ and inhibition zones were observed. A positive zone of inhibition (sensitive, 15 mm) is usually in case of pneumococci and no inhibition (resistant) is usually seen in viridans group or other alpha-hemolytic streptococci. The type of hemolysis was also noted during this assay. Finally, serotyping was performed using standard Quellung’s (Neufeld) reaction(95) towards capsular polysaccharide as described(96).

### Antibiotic Susceptibility Tests

Antibiograms and Minimum Inhibitory Concentration (MIC) studies were performed at the Institute of Infectious Diseases, University of Bern, Switzerland, according to the criteria established by European Committee on Antimicrobial Susceptibility Testing ([EUCAST](https://www.eucast.org/clinical_breakpoints))(96). Bacteria were grown on Muller-Hinton agar for Fastidious organisms (MH supplemented with 5% defibrinated horse blood and 20 mg/l β-NAD). Antibiograms were performed using disk diffusion method (disk content in μg), observing zone of inhibition (diameter in mm). MIC determination was performed using Etest® strips (Biomérieux, France, described in μg/ml). The values were tallied with the EUCAST v13.0 clinical breakpoint tables for interpretation of results. Macrolide-inducible resistance to clindamycin test (D-test) to test for assessing macrolide-lincosamide-streptogramin B (MLS_B_) resistance, was performed by placing Erythromycin and Clindamycin disks 12-20 mm apart (edge to edge). Appearance of antagonism (the D phenomenon) was observed to detect any inducible clindamycin resistance.

### Biochemical identification of *Streptococcaceae*

Streptococcal identification was carried out by using API® 20 Strep panel (Biomérieux, France), a standardized system combining 20 biochemical tests, according to manufacturer’s instructions. Briefly, all isolates were first grown as previously described (Methods: Bacterial growth and media). Bacteria were collected using a sterile cotton swab and mixed in 2ml API® Suspension Medium to achieve turbidity more than McFarland standard 4 before distributing into the given cupules in the panel strips and incubating according to manufacturer’s instructions. In this case, both 4- and 24-hour readings were performed as some isolates tend to exhibit delayed effects.

## Functional annotation and analysis

### Metabolic profiling and macromolecular structures prediction

Predictions of key metabolic pathways and macromolecular structures were performed by using a custom genome profiler as previously described(28). A set of rules were applied to conclude the presence and completeness of predicted pathways. These can be found in the scripts defining the rules at https://github.com/gsartonl/Publication_Sarton-Loheac_2022. Metabolic predictions were performed using GapMind(30) for amino acid biosynthesis and dbCAN(97) for Carbohydrate Active enzymes(98) ([CAZyme](http://www.cazy.org/)) analysis (Dataset S11). For secretion systems and other macromolecular structures, MacSyFinder(99) was used. The results were visualized along as heatmap depicting system completeness (%) along with a single copy core genes-based phylogeny using [iTOL](https://itol.embl.de/)(88) v6.8.1 webtool.

### COG categories

eggNOG (evolutionary genealogy of genes: nonsupervised orthologous groups) was used to predict the functional annotation of genes. Representative genes from six individual Streptococcal species and the 977 shared genes among the six isolates obtained from the OrthoVenn2 tool were used. Shared genes from any one of the files containing the protein FASTA sequences of the 6 isolates was performed using *Append_concatenate_extract_grep_Linux_log.md and Extraction_of_headers_fasta_sequences_and_splitted_files.md*. Protein (up to 100,000) sequences from the genomes of individual species along with those shared 977 extracted sequences was uploaded separately to the eggnog-mapper v2 web tool(27). The individual output files (.xlsx) were downloaded (Dataset S10). The COGs assigned to different individual proteins was used to calculate frequencies in each of the isolates using *COG_analysis_Rscripts.md*. This was visualized as heatmap along with a single copy core genes-based phylogeny using [iTOL](https://itol.embl.de/)(88) v6.8.1 webtool.

### Anti-microbial and Virulence genes analysis

Genes coding for antimicrobial resistance (AMR) and known virulence factors were detected by *Antimicrobial_resistance_virulence_gene_search.md* that uses [ABRicatetool](https://github.com/tseemann/abricate) (v 1.0.1). The databases used were MEGARes(46) and VFDB(51) for AMR and virulence factors respectively. Genomic DNA FASTA sequences of six Streptococcal species along with the 47 reference genomes were used as input files. The output text file (.csv) was summarized using abrica– -- summary command, which included genome ID, gene name, percent coverage and number of genes found. The results were visualized along as heatmap depicting percentage (%) coverage along with a single copy core genes-based phylogeny using [iTOL](https://itol.embl.de/)(88) v6.8.1 webtool.

### Screening of Streptococcal capsule in silico

For capsule analysis, the prototypical *Streptococcus pneumonaie* capsule locus was used as the query and BLAST command line tool was used to search for matching proteins in the isolates and reference genomes. Protein FASTA sequences of capsule genes i.e., 17 proteins encoded by *cps*A-T were extracted from the D39V genome and saved as a single FASTA file (Dataset S13). For creating a custom BLAST database, all proteins from 6 isolates and 47 reference genomes were combined to create the Pan-Strep database containing 80,364 protein sequences. The database was created under the ‘*ncbi-datasets*’ environment in command line using the ‘*makeblastdb*’ function. Full script and analysis steps are described in ‘*NCBI datasets and BLAST.md*’. The FASTA file containing query capsule proteins was used to perform BLASTP 2.6.0 search and the output was obtained in a tabular format. A set of rules were decided for the protein to be considered as a significant match. These were: (i) E-value cut-off at 0.0, (ii) Bit score > 200 and (iii) identity threshold of > 50%. The protein IDs of the resulting matches used to extract the FASTA sequences using *Append_concatenate_extract_grep_Linux_log.md* and *Extraction_of_headers_fasta_sequences_and_splitted_files.md.* The capsule gene sequences were rearranged according to bacterial genomes and were concatenated followed by *Alignment and phylogeny.md* (50% sites stripped, RAxML model ‘PROTGAMMAAUTO’, 100 rapid bootstrapping). The results were visualized along as barchart showing distribution and number of capsule genes found in each genome along phylogeny using [iTOL](https://itol.embl.de/)(88) v6.8.1 webtool. Gene plots were constructed using pyGenomeViz(100) to visualize the genomic arrangement and synteny of lung Streptococcal isolates in comparison to the prototypical capsule locus of *Streptococcous pneumonaie* D39.

### Pneumolysin analysis

Pneumolysin protein encoded by the *ply* gene, from the prototypical gene sequence from *Streptococcus pneumonaie* D39 was used as a query to perform BLASTP 2.6.0 search against the custom Pan-Strep database. The hits found were then extracted from the respective genomes and followed by *Alignment and phylogeny.md* (MUSCLE(101), RAxML model ‘PROTGAMMAAUTO’, 100 rapid bootstrapping) (Dataset S12). For visualization, a heatmap with distance matrix was constructed in R and gene plot was made using pyGenomeViz.

### Gene plots and visualization of genomic features

For visualizing sequence similarity and comparison of gene arrangements between multiple genomes pyGenomeViz-MMSeqs v0.3.2 tool was to plot genomic features. This tool was used in the conda environment with default parameters using Genbank files (.gb or .gbk) as input files. Output data contained the reciprocal best hits file (.tsv) and the syntenic plots between the genomes. In synteny plots, pairwise sequence similarity can be observed between genomes or coding sequences. The position of the genes can be compared using the genomic coordinates. This synteny analysis also informs about reciprocal mapping to identify regions of similarity and orthologous genes for understanding the evolutionary relationships.

## Acknowledgements

Funding for this work came from several agencies for different contributors to this work, which are as follows. Sudip Das was funded by Marie Sklodowska-Curie Individual Fellowship (“HUMANITY”, Grant no. 800301) and an Interdisciplinary grant from the Faculty of Biology and Medicine of the University of Lausanne (awarded to Philipp Engel as supervisor, Grant no. 26075716). Philipp Engel is funded by an ERC StG (“MicroBeeOme”, Grant No. 714804), two Swiss National Science Foundation grants (Grant no. 31003A_160345 and 31003A_179487). Germán-Bonilla Rosso was funded by HFSP Young Investigator grant (awarded to Philipp Engel as supervisor, Grant no. RGY0077/2016). We thank Prof. Dr. Jan-Willem Veening and his team at the Department of Fundamental Microbiology, University of Lausanne, Lausanne, Switzerland for providing the bacterial strains *S. pneumonie D39* and *S. mitis NCTC* 12261.

## References

1. Charlson ES, Bittinger K, Haas AR, Fitzgerald AS, Frank I, Yadav A, Bushman FD, Collman RG. 2011. Topographical continuity of bacterial populations in the healthy human respiratory tract. Am J Respir Crit Care Med 10.1164/rccm.201104-0655OC.

2. Das S, Bernasconi E, Koutsokera A, Wurlod D-A, Tripathi V, Bonilla-Rosso G, Aubert J-D, Derkenne M-F, Mercier L, Pattaroni C, Rapin A, von Garnier C, Marsland BJ, Engel P, Nicod LP. 2021. A prevalent and culturable microbiota links ecological balance to clinical stability of the human lung after transplantation. Nat Commun 12:2126-undefined.

3. Pattaroni C, Watzenboeck ML, Schneidegger S, Kieser S, Wong NC, Bernasconi E, Pernot J, Mercier L, Knapp S, Nicod LP, Marsland CP, Roth-Kleiner M, Marsland BJ. 2018. Early-Life Formation of the Microbial and Immunological Environment of the Human Airways. Cell Host Microbe 24:857–865.e4.

4. Bernasconi E, Pattaroni C, Koutsokera A, Pison C, Kessler R, Benden C, Soccal PM, Magnan A, Aubert J-D, Marsland BJ, Nicod LP. 2016. Airway Microbiota Determines Innate Cell Inflammatory or Tissue Remodeling Profiles in Lung Transplantation. Am J Respir Crit Care Med 194:1252–1263.

5. Venkataraman A, Bassis CM, Beck JM, Young VB, Curtis JL, Huffnagle GB, Schmidt TM. 2015. Application of a Neutral Community Model To Assess Structuring of the Human Lung Microbiome. mBio 6.

6. Marsland BJ, Gollwitzer ES. 2014. Host–microorganism interactions in lung diseases. Nat Rev Immunol 14:827–835.

7. Man WH, De Steenhuijsen Piters WAA, Bogaert D. 2017. The microbiota of the respiratory tract: Gatekeeper to respiratory health. Nat Rev Microbiol 10.1038/nrmicro.2017.14.

8. Dickson RP, Huffnagle GB. 2015. The Lung Microbiome: New Principles for Respiratory Bacteriology in Health and Disease. PLoS Pathog 11:e1004923.

9. Bassis CM, Erb-Downward JR, Dickson RP, Freeman CM, Schmidt TM, Young VB, Beck JM, Curtis JL, Huffnagle GB. 2015. Analysis of the Upper Respiratory Tract Microbiotas as the Source of the Lung and Gastric Microbiotas in Healthy Individuals. mBio 6.

10. Dickson RP, Erb-Downward JR, Martinez FJ, Huffnagle GB. 2016. The Microbiome and the Respiratory Tract. Annu Rev Physiol 78:481–504.

11. Dickson RP, Erb-Downward JR, Freeman CM, McCloskey L, Beck JM, Huffnagle GB, Curtis JL. 2015. Spatial Variation in the Healthy Human Lung Microbiome and the Adapted Island Model of Lung Biogeography. Ann Am Thorac Soc 12:821–830.

12. Lloyd CM, Marsland BJ. 2017. Lung Homeostasis: Influence of Age, Microbes, and the Immune System. Immunity 46:549–561.

13. Whelan FJ, Heirali AA, Rossi L, Rabin HR, Parkins MD, Surette MG. 2017. Longitudinal sampling of the lung microbiota in individuals with cystic fibrosis. PLoS One 10.1371/journal.pone.0172811.

14. Hilty M, Burke C, Pedro H, Cardenas P, Bush A, Bossley C, Davies J, Ervine A, Poulter L, Pachter L, Moffatt MF, Cookson WOC. 2010. Disordered microbial communities in asthmatic airways. PLoS One 10.1371/journal.pone.0008578.

15. Mika M, Nita I, Morf L, Qi W, Beyeler S, Bernasconi E, Marsland BJ, Ott SR, von Garnier C, Hilty M. 2018. Microbial and host immune factors as drivers of COPD. ERJ Open Res 4:00015–02018.

16. Woo TE, Lim R, Heirali AA, Acosta N, Rabin HR, Mody CH, Somayaji R, Surette MG, Sibley CD, Storey DG, Parkins MD. 2019. A longitudinal characterization of the Non-Cystic Fibrosis Bronchiectasis airway microbiome. Sci Rep 10.1038/s41598-019-42862-y.

17. Afrizal A, Hitch TCA, Viehof A, Treichel N, Riedel T, Abt B, Buhl EM, Kohlheyer D, Overmann J, Clavel T. 2022. Anaerobic single-cell dispensing facilitates the cultivation of human gut bacteria. Environ Microbiol 10.1111/1462-2920.15935.

18. Das S, Bernasconi E, Koutsokera A, Wurlod D-A, Tripathi V, Bonilla-Rosso G, Aubert J-D, Derkenne M-F, Mercier L, Pattaroni C, Rapin A, von Garnier C, Marsland BJ, Engel P, Nicod LP. 2021. A prevalent and culturable microbiota links ecological balance to clinical stability of the human lung after transplantation. Nat Commun 12.

19. Auch AF, von Jan M, Klenk HP, Göker M. 2010. Digital DNA-DNA hybridization for microbial species delineation by means of genome-to-genome sequence comparison. Stand Genomic Sci 2:117.

20. Meier-Kolthoff JP, Göker M. 2019. TYGS is an automated high-throughput platform for state-of-the-art genome-based taxonomy. Nat Commun 10.

21. Jain C, Rodriguez-R LM, Phillippy AM, Konstantinidis KT, Aluru S. 2018. High throughput ANI analysis of 90K prokaryotic genomes reveals clear species boundaries. Nat Commun 9:5114.

22. Beighton D, Hardie JM, Whiley RA. 1991. A scheme for the identification of viridans streptococci. J Med Microbiol 35:367–372.

23. Kawamura Y, Hou XG, Sultana F, Miura H, Ezaki T. 1995. Determination of 16S rRNA sequences of Streptococcus mitis and Streptococcus gordonii and phylogenetic relationships among members of the genus Streptococcus. Int J Syst Bacteriol 45:406– 408.

24. Delorme C, Abraham AL, Renault P, Guédon E. 2015. Genomics of Streptococcus salivarius, a major human commensal. Infection, Genetics and Evolution 33:381–392.

25. Emms DM, Kelly S. 2019. OrthoFinder: Phylogenetic orthology inference for comparative genomics. Genome Biol 20:1–14.

26. Xu L, Dong Z, Fang L, Luo Y, Wei Z, Guo H, Zhang G, Gu YQ, Coleman-Derr D, Xia Q, Wang Y. 2019. OrthoVenn2: a web server for whole-genome comparison and annotation of orthologous clusters across multiple species. Nucleic Acids Res 47:W52.

27. Cantalapiedra CP, Hern̗andez-Plaza A, Letunic I, Bork P, Huerta-Cepas J. 2021. eggNOG-mapper v2: Functional Annotation, Orthology Assignments, and Domain Prediction at the Metagenomic Scale. Mol Biol Evol 38:5825–5829.

28. Sarton-Lohéac G, Nunes da Silva CG, Mazel F, Baud G, de Bakker V, Das S, El Chazli Y, Ellegaard K, Garcia-Garcera M, Glover N, Liberti J, Nacif Marçal L, Prasad A, Somerville V, Bonilla-Rosso G, Engel P. 2023. Deep Divergence and Genomic Diversification of Gut Symbionts of Neotropical Stingless Bees. mBio 14:e0353822.

29. Price MN, Deutschbauer AM, Arkin AP. 2022. Filling gaps in bacterial catabolic pathways with computation and high-throughput genetics. PLoS Genet 18:e1010156.

30. Price MN, Deutschbauer AM, Arkin AP. 2020. GapMind: Automated Annotation of Amino Acid Biosynthesis. mSystems 5.

31. Zheng J, Ge Q, Yan Y, Zhang X, Huang L, Yin Y. 2023. dbCAN3: automated carbohydrate-active enzyme and substrate annotation. Nucleic Acids Res 51:W115– W121.

32. Abby SS, Néron B, Ménager H, Touchon M, Rocha EPC. 2014. MacSyFinder: A Program to Mine Genomes for Molecular Systems with an Application to CRISPR-Cas Systems. PLoS One 9:e110726.

33. Cantarel BI, Coutinho PM, Rancurel C, Bernard T, Lombard V, Henrissat B. 2009. The Carbohydrate-Active EnZymes database (CAZy): An expert resource for glycogenomics. Nucleic Acids Res 37.

34. Peterson SN, Sung CK, Cline R, Desai B V., Snesrud EC, Luo P, Walling J, Li H, Mintz M, Tsegaye G, Burr PC, Do Y, Ahn S, Gilbert J, Fleischmann RD, Morrison DA. 2004. Identification of competence pheromone responsive genes in Streptococcus pneumoniae by use of DNA microarrays. Mol Microbiol 51:1051–1070.

35. Whatmore AM, Barcus VA, Dowson CG. 1999. Genetic Diversity of the Streptococcal Competence (*com*) Gene Locus. J Bacteriol 181:3144–3154.

36. Luo P, Morrison DA. 2003. Transient association of an alternative sigma factor, ComX, with RNA polymerase during the period of competence for genetic transformation in Streptococcus pneumoniae. J Bacteriol 185:349–358.

37. Provvedi R, Dubnau D. 1999. ComEA is a DNA receptor for transformation of competent Bacillus subtilis. Mol Microbiol 31:271–280.

38. Diallo A, Foster HR, Gromek KA, Perry TN, Dujeancourt A, Krasteva P V., Gubellini F, Falbel TG, Burton BM, Fronzes R. 2017. Bacterial transformation: ComFA is a DNA-dependent ATPase that forms complexes with ComFC and DprA. Mol Microbiol 105:741–754.

39. Merritt J, Qi F, Shi W. 2005. A unique nine-gene comY operon in Streptococcus mutans. Microbiology (Reading) 151:157–166.

40. Haijema BJ, Hahn J, Haynes J, Dubnau D. 2001. A ComGA-dependent checkpoint limits growth during the escape from competence. Mol Microbiol 40:52–64.

41. Guglielmini J, Néron B, Abby SS, Garcillán-Barcia MP, La Cruz DF, Rocha EPC. 2014. Key components of the eight classes of type IV secretion systems involved in bacterial conjugation or protein secretion. Nucleic Acids Res 42:5715.

42. Atmakuri K, Cascales E, Christie PJ. 2004. Energetic components VirD4, VirB11 and VirB4 mediate early DNA transfer reactions required for bacterial type IV secretion. Mol Microbiol 54:1199–1211.

43. Kirn TJ, Bose N, Taylor RK. 2003. Secretion of a soluble colonization factor by the TCP type 4 pilus biogenesis pathway in Vibrio cholerae. Mol Microbiol 49:81–92.

44. Waksman G. 2019. From conjugation to T4S systems in Gram-negative bacteria: a mechanistic biology perspective. EMBO Rep 20:e47012.

45. Seemann T. GitHub - tseemann/abricate: :mag_right: Mass screening of contigs for antimicrobial and virulence genes. https://github.com/tseemann/abricate. Retrieved 10 November 2023.

46. Doster E, Lakin SM, Dean CJ, Wolfe C, Young JG, Boucher C, Belk KE, Noyes NR, Morley PS. 2020. MEGARes 2.0: a database for classification of antimicrobial drug, biocide and metal resistance determinants in metagenomic sequence data. Nucleic Acids Res 48:D561–D569.

47. Daly MM, Doktor S, Flamm R, Shortridge D. 2004. Characterization and Prevalence of MefA, MefE, and the Associated *msr* (D) Gene in *Streptococcus pneumoniae* Clinical Isolates. J Clin Microbiol 42:3570–3574.

48. Clancy J, Petitpas J, Dib-Hajj F, Yuan W, Cronan M, Kamath A V., Bergeron J, Retsema JA. 1996. Molecular cloning and functional analysis of a novel macrolide-resistance determinant, mefA, from Streptococcus pyogenes. Mol Microbiol 22:867– 879.

49. Leclercq R, Courvalin P. 1991. Bacterial resistance to macrolide, lincosamide, and streptogramin antibiotics by target modification. Antimicrob Agents Chemother 35:1267–1272.

50. Warburton PJ, Ciric L, Lerner A, Seville LA, Roberts AP, Mullany P, Allan E. 2013. TetAB(46), a predicted heterodimeric ABC transporter conferring tetracycline resistance in Streptococcus australis isolated from the oral cavity. Journal of Antimicrobial Chemotherapy 68:17.

51. Chen L, Yang J, Yu J, Yao Z, Sun L, Shen Y, Jin Q. 2005. VFDB: a reference database for bacterial virulence factors. Nucleic Acids Res 33.

52. Hu DK, Wang DG, Liu Y, Liu CB, Yu LH, Qu Y, Luo XH, Yang JH, Yu J, Zhang J, Li XY. 2013. Roles of virulence genes (PsaA and CpsA) on the invasion of Streptococcus pneumoniae into blood system. Eur J Med Res 18:1–6.

53. Hu Y, Park N, Seo KS, Park JY, Somarathne RP, Olivier AK, Fitzkee NC, Thornton JA. 2021. Pneumococcal surface adhesion A protein (PsaA) interacts with human Annexin A2 on airway epithelial cells. Virulence 12:1841–1854.

54. Carvalho MDGS, Tondella ML, McCaustland K, Weidlich L, McGee L, Mayer LW, Steigerwalt A, Whaley M, Facklam RR, Fields B, Carlone G, Ades EW, Dagan R, Sampson JS. 2007. Evaluation and Improvement of Real-Time PCR Assays Targeting lytA, ply, and psaA Genes for Detection of Pneumococcal DNA. J Clin Microbiol 45:2460–2466.

55. Skov Sørensen UB, Yao K, Yang Y, Tettelin H, Kilian M. 2016. Capsular polysaccharide expression in commensal Streptococcus species: Genetic and antigenic similarities to Streptococcus pneumoniae. mBio 7.

56. Agapov VS, Smirenskaia T V, Komnova ZD. 1987. [Clinico-morphological characteristics of periradicular cysts bordering on the maxillary sinus]. Stomatologiia (Mosk) 66:11–13.

57. Guerin ME, Kordulakova J, Schaeffer F, Svetlikova Z, Buschiazzo A, Giganti D, Gicquel B, Mikusova K, Jackson M, Alzari PM. 2007. Molecular recognition and interfacial catalysis by the essential phosphatidylinositol mannosyltransferase PimA from mycobacteria. J Biol Chem 282:20705–20714.

58. Slager J, Aprianto R, Veening JW. 2018. Deep genome annotation of the opportunistic human pathogen Streptococcus pneumoniae D39. Nucleic Acids Res 46:9971–9989.

59. Mohanty P, Patel A, Bhardwaj AK. 2012. Role of H- and D- MATE-type transporters from multidrug resistant clinical isolates of Vibrio fluvialis in conferring fluoroquinolone resistance. PLoS One 7.

60. Smith HE, Damman M, van der Velde J, Wagenaar F, Wisselink HJ, Stockhofe-Zurwieden N, Smits MA. 1999. Identification and Characterization of the *cps* Locus of *Streptococcus suis* Serotype 2: the Capsule Protects against Phagocytosis and Is an Important Virulence Factor. Infect Immun 67:1750–1756.

61. Escapa IF, Chen T, Huang Y, Gajare P, Dewhirst FE, Lemon KP. 2018. New Insights into Human Nostril Microbiome from the Expanded Human Oral Microbiome Database (eHOMD): a Resource for the Microbiome of the Human Aerodigestive Tract. mSystems 3.

62. Segal LN, Alekseyenko A V., Clemente JC, Kulkarni R, Wu B, Chen H, Berger KI, Goldring RM, Rom WN, Blaser MJ, Weiden MD. 2013. Enrichment of lung microbiome with supraglottic taxa is associated with increased pulmonary inflammation. Microbiome 1:19.

63. Erb-Downward JR, Thompson DL, Han MK, Freeman CM, McCloskey L, Schmidt LA, Young VB, Toews GB, Curtis JL, Sundaram B, Martinez FJ, Huffnagle GB. 2011. Analysis of the lung microbiome in the “healthy” smoker and in COPD. PLoS One 10.1371/journal.pone.0016384.

64. Charlson ES, Diamond JM, Bittinger K, Fitzgerald AS, Yadav A, Haas AR, Bushman FD, Collman RG. 2012. Lung-enriched Organisms and Aberrant Bacterial and Fungal Respiratory Microbiota after Lung Transplant. Am J Respir Crit Care Med 186:536– 545.

65. Borewicz K, Pragman AA, Kim HB, Hertz M, Wendt C, Isaacson RE. 2013. Longitudinal analysis of the lung microbiome in lung transplantation. FEMS Microbiol Lett 10.1111/1574-6968.12053.

66. Sharma NS, Vestal G, Wille K, Patel KN, Cheng F, Tipparaju S, Tousif S, Banday MM, Xu X, Wilson L, Nair VS, Morrow C, Hayes D, Seyfang A, Barnes S, Deshane JS, Gaggar A. 2020. Differences in airway microbiome and metabolome of single lung transplant recipients. Respir Res 21:1–12.

67. Doern CD, Carey-Ann BD. 2010. It’s Not Easy Being Green: the Viridans Group Streptococci, with a Focus on Pediatric Clinical Manifestations. J Clin Microbiol 48:3829–3835.

68. Ruoff KL. 1991. Nutritionally Variant Streptococci. Clin Microbiol Rev 4:184–190.

69. Kim JN, Ahn SJ, Burne RA. 2015. Genetics and Physiology of Acetate Metabolism by the Pta-Ack Pathway of Streptococcus mutans. Appl Environ Microbiol 81:5015.

70. Garnett JP, Baker EH, Baines DL. 2012. Sweet talk: insights into the nature and importance of glucose transport in lung epithelium. European Respiratory Journal 40:1269–1276.

71. Kwong WK, Zheng H, Moran NA. 2018. Erratum to: Convergent evolution of a modified, acetate-driven TCA cycle in bacteria (Nature Microbiology, (2017), 2, (17067), 10.1038/nmicrobiol.2017.67). Nat Microbiol 3:960.

72. Sawers RG, Clark DP. 2004. Fermentative Pyruvate and Acetyl-Coenzyme A Metabolism. EcoSal Plus 1.

73. Tagaino R, Washio J, Abiko Y, Tanda N, Sasaki K, Takahashi N. 2019. Metabolic property of acetaldehyde production from ethanol and glucose by oral Streptococcus and Neisseria. Scientific Reports 2019 9:1 9:1–8.

74. Deleu S, Machiels K, Raes J, Verbeke K, Vermeire S. 2021. Short chain fatty acids and its producing organisms: An overlooked therapy for IBD? EBioMedicine 66:103293.

75. Antunes KH, Fachi JL, de Paula R, da Silva EF, Pral LP, dos Santos AÁ, Dias GBM, Vargas JE, Puga R, Mayer FQ, Maito F, Zárate-Bladés CR, Ajami NJ, Sant’Ana MR, Candreva T, Rodrigues HG, Schmiele M, Silva Clerici MTP, Proença-Modena JL, Vieira AT, Mackay CR, Mansur D, Caballero MT, Marzec J, Li J, Wang X, Bell D, Polack FP, Kleeberger SR, Stein RT, Vinolo MAR, de Souza APD. 2019. Microbiota-derived acetate protects against respiratory syncytial virus infection through a GPR43-type 1 interferon response. Nat Commun 10:3273.

76. Machado MG, Patente TA, Rouillé Y, Heumel S, Melo EM, Deruyter L, Pourcet B, Sencio V, Teixeira MM, Trottein F. 2022. Acetate Improves the Killing of Streptococcus pneumoniae by Alveolar Macrophages via NLRP3 Inflammasome and Glycolysis-HIF-1α Axis. Front Immunol 13.

77. Håvarstein LS, Coomaraswamy G, Morrison DA. 1995. An unmodified heptadecapeptide pheromone induces competence for genetic transformation in Streptococcus pneumoniae. Proc Natl Acad Sci U S A 92:11140.

78. Haenni M, Lupo A, Madec J-Y. 2018. Antimicrobial Resistance in Streptococcus spp . Microbiol Spectr 6.

79. Gibson PS, Veening J-W. 2023. Gaps in the wall: understanding cell wall biology to tackle amoxicillin resistance in Streptococcus pneumoniae. Curr Opin Microbiol 72:102261.

80. Skov Sørensen UB, Yao K, Yang Y, Tettelin H, Kilian M. 2016. Capsular Polysaccharide Expression in Commensal *Streptococcus* Species: Genetic and Antigenic Similarities to Streptococcus pneumoniae. mBio 7.

81. Chang B, Morita M, Nariai A, Kasahara K, Kakutani A, Ogawa M, Ohnishi M, Oishi K. 2022. Invasive Streptococcus oralis Expressing Serotype 3 Pneumococcal Capsule, Japan - Volume 28, Number 8—August 2022 - Emerging Infectious Diseases journal - CDC. Emerg Infect Dis 28:1720–1722.

82. s-andrews/FastQC: A quality control analysis tool for high throughput sequencing data. https://github.com/s-andrews/FastQC. Retrieved 13 July 2023.

83. Bolger AM, Lohse M, Usadel B. 2014. Trimmomatic: a flexible trimmer for Illumina sequence data. Bioinformatics 30:2114–2120.

84. Bankevich A, Nurk S, Antipov D, Gurevich AA, Dvorkin M, Kulikov AS, Lesin VM, Nikolenko SI, Pham S, Prjibelski AD, Pyshkin A V., Sirotkin A V., Vyahhi N, Tesler G, Alekseyev MA, Pevzner PA. 2012. SPAdes: a new genome assembly algorithm and its applications to single-cell sequencing. J Comput Biol 19:455–477.

85. Mikheenko A, Saveliev V, Gurevich A. 2016. MetaQUAST: evaluation of metagenome assemblies. Bioinformatics 32:1088–1090.

86. Parks DH, Imelfort M, Skennerton CT, Hugenholtz P, Tyson GW. 2015. CheckM: assessing the quality of microbial genomes recovered from isolates, single cells, and metagenomes. Genome Res 25:1043–1055.

87. Seemann T. 2014. Prokka: rapid prokaryotic genome annotation. Bioinformatics 30:2068–2069.

88. Letunic I, Bork P. 2007. Interactive Tree Of Life (iTOL): an online tool for phylogenetic tree display and annotation. Bioinformatics 23:127–128.

89. Jain C, Rodriguez-R LM, Phillippy AM, Konstantinidis KT, Aluru S. 2018. High throughput ANI analysis of 90K prokaryotic genomes reveals clear species boundaries. Nature Communications 2018 9:1 9:1–8.

90. aniclustermap · PyPI. https://pypi.org/project/aniclustermap/. Retrieved 15 June 2023.

91. Katoh K, Misawa K, Kuma KI, Miyata T. 2002. MAFFT: a novel method for rapid multiple sequence alignment based on fast Fourier transform. Nucleic Acids Res 30:3059–3066.

92. Price MN, Dehal PS, Arkin AP. 2009. FastTree: Computing Large Minimum Evolution Trees with Profiles instead of a Distance Matrix. Mol Biol Evol 26:1641– 1650.

93. Stamatakis A. 2006. RAxML-VI-HPC: maximum likelihood-based phylogenetic analyses with thousands of taxa and mixed models. Bioinformatics 22:2688–2690.

94. Mostacci N, Wüthrich TM, Siegwald L, Kieser S, Steinberg R, Sakwinska O, Latzin P, Korten I, Hilty M. 2023. Informed interpretation of metagenomic data by StrainPhlAn enables strain retention analyses of the upper airway microbiome. mSystems 8:e0072423.

95. Neufeld F. 1902. Ueber die Agglutination der Pneumokokken und über die Theorieen der Agglutination. Zeitschrift für Hygiene und Infektionskrankheiten 1902 40:1 40:54–72.

96. Agapov VS, Smirenskaia T V, Komnova ZD. 1987. [Clinico-morphological characteristics of periradicular cysts bordering on the maxillary sinus]. Stomatologiia (Mosk) 66:11–13.

97. Yin Y, Mao X, Yang J, Chen X, Mao F, Xu Y. 2012. dbCAN: a web resource for automated carbohydrate-active enzyme annotation. Nucleic Acids Res 40:W445– W451.

98. Drula E, Garron ML, Dogan S, Lombard V, Henrissat B, Terrapon N. 2022. The carbohydrate-active enzyme database: functions and literature. Nucleic Acids Res 50:D571–D577.

99. Abby SS, Néron B, Ménager H, Touchon M, Rocha EPC. 2014. MacSyFinder: A Program to Mine Genomes for Molecular Systems with an Application to CRISPR-Cas Systems. PLoS One 9:e110726.

100. Shimoyama Y. 2022. pyGenomeViz: A genome visualization python package for comparative genomics. https://github.com/moshi4/pyGenomeViz. Retrieved 14 December 2023.

101. Edgar RC. 2004. MUSCLE: a multiple sequence alignment method with reduced time and space complexity. BMC Bioinformatics 5:113.

