## Supplementary Information for "Comparative genomic analysis reveals novel phylogenetically intermediate Streptococci with high phenotypic diversity in the human distal lung microbiota"

### Supplementary Figures

**Figure S1**

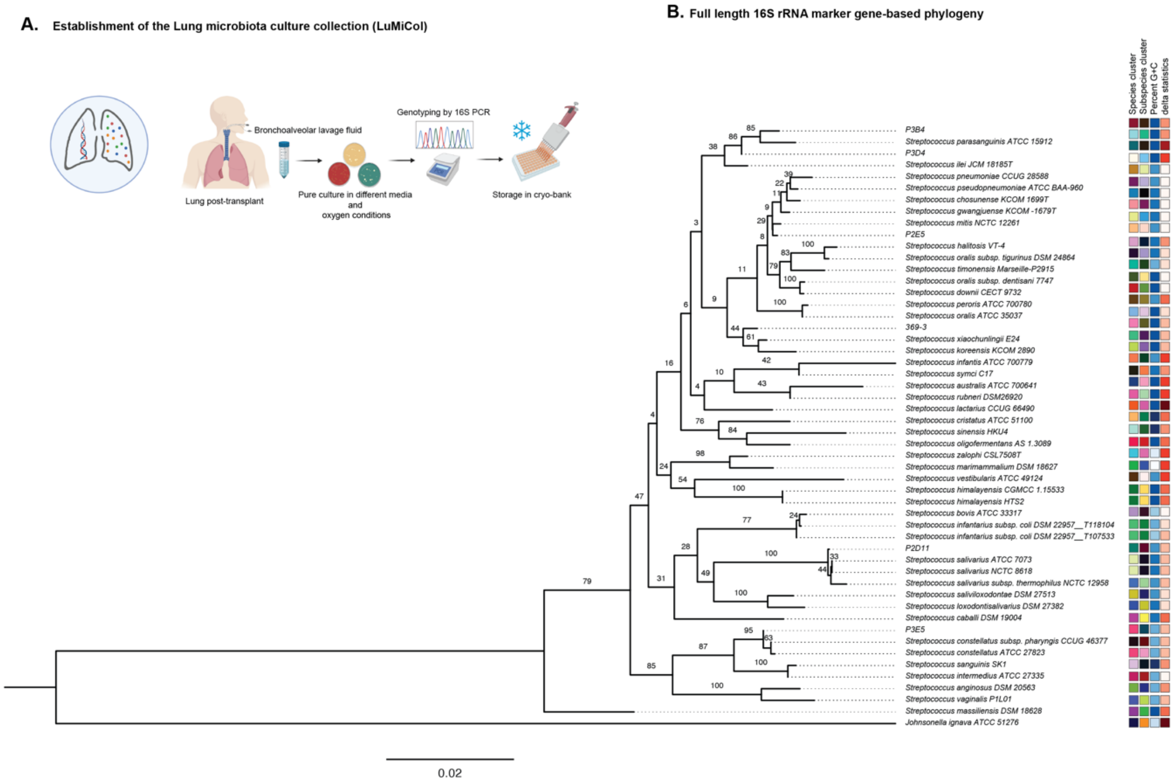


**Figure S1. A.** Workflow showing establishment of the human Lung microbiota culture collection (LuMiCol)**. It is** an open-source bacterial biobank created by large scale bacterial culturing of Bronchoalveolar lavage fluid samples post lung transplant at varied culture conditions including different media and oxygen availability. Genotyping was performed by amplification of 16S rRNA gene after pure culture of single colonies. Confirmed isolates were grown and stored in 96-well plates in cryo-storage (-80°C). **B.** Comparison of human distal lung streptococci to closely related type strains in the TYGS database. Full-length 16S rRNA gene BLAST Distance Phylogeny (GBDP) using FASTME, where colored boxes represent species and subspecies clusters, blue-colored gradient boxes represent GC content (%) and brown-coloured gradient represent delta statistics.

**Figure S2**

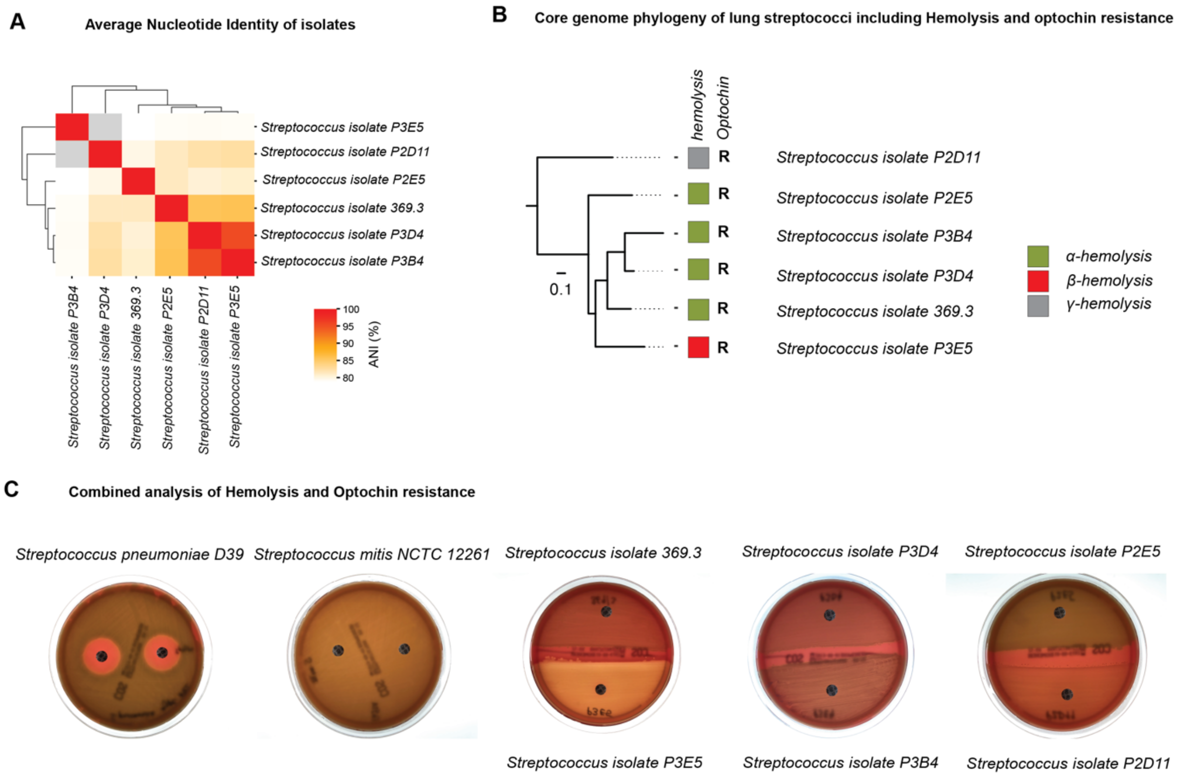


**Figure S2. A**. Heatmap showing pairwise Average Nucleotide Identity (ANI %) between human distal lung streptococci. ANI values > 80% were colored as grey in the heatmap. **B.** Optochin resistance and alpha hemolysis of lung streptococcal isolates depicted according to maximum-likelihood phylogeny computed computed by FastTree on concatenated amino acid sequences of 957 single-copy core proteins. Hemolysis type was depicted by colored boxes. **C.** Images of combined tests for optochin resistance (disk diffusion) and hemolysis performed on lung human lung isolates along with the controls: *S. pneumoniae* D39 and *S. mitis* NCTC 12261.

**Figure S3**

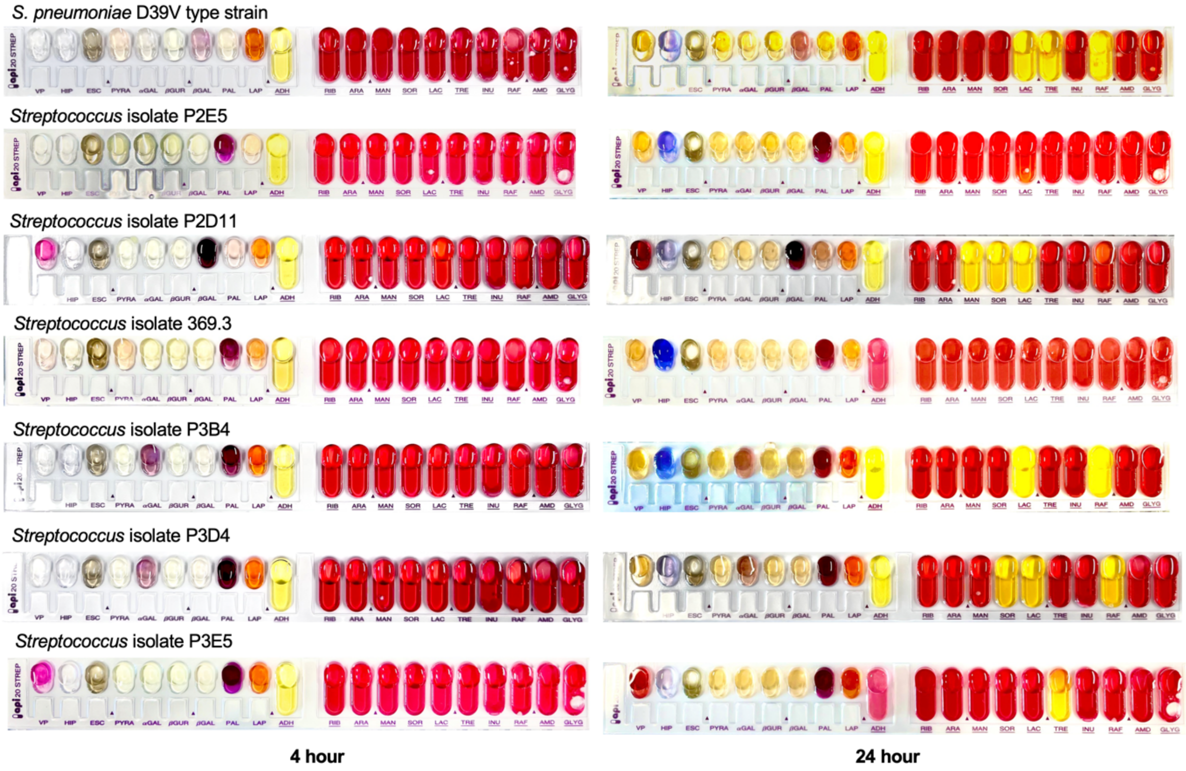


**Figure S3.** StrepAPI® 20 panels for biochemical characterization of human distal lung streptococcal isolates along *S. pneumoniae* D39V as control. Test results were observed at 4 hr (left) and 24 hour (right) post inoculation.

### Figure S4

**A** **Total number of CAZyme families found in Pan-Strep and lung streptococci**

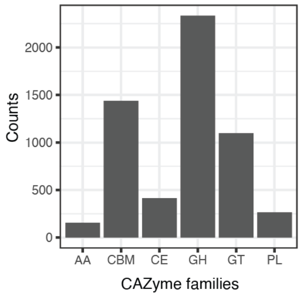


**B** **Summary of Glycosyl Hydrolase (GH) types present in Pan-Strep and lung streptococci**

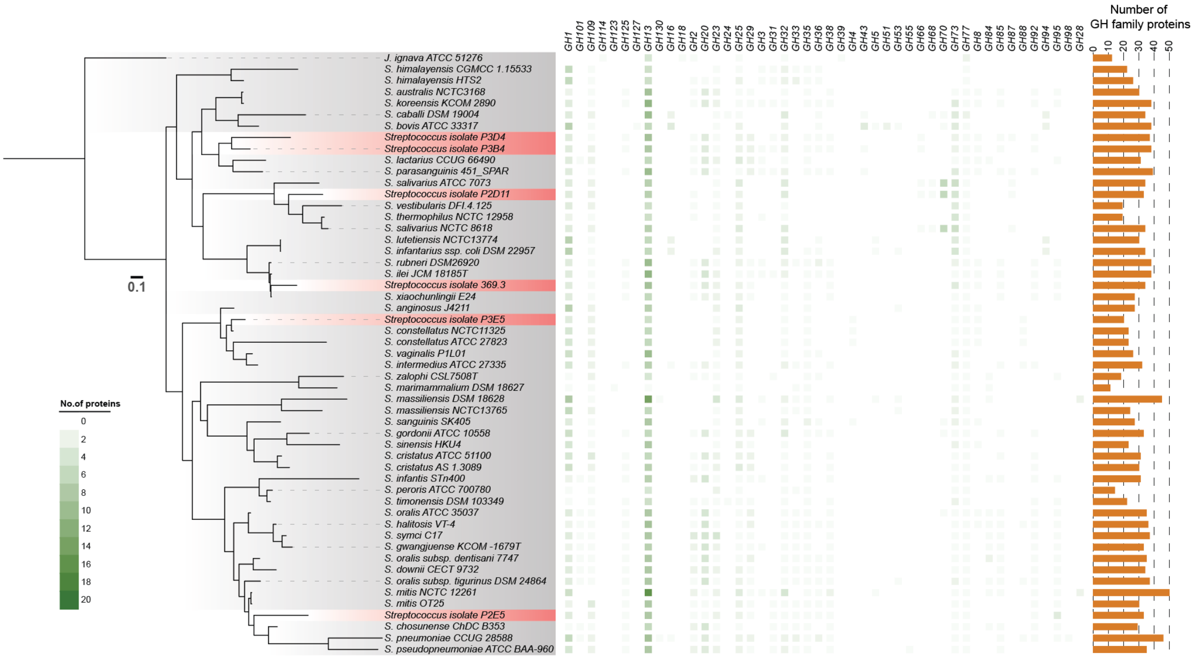


**Figure S4. A.** Bar chart showing the total number of Carbohydrate-active enzyme (CAZyme) families (y-axis) grouped according to family type (AA, CBM, CE, GH, GT, PL, x-axis) found in all genomes (Pan-Strep database and lung isolates). **B.** Heatmap showing number of proteins in each Glycosyl Hydrolase (GH) family type depicted according to maximum-likelihood phylogeny computed by FastTree on concatenated amino acid sequences of 315 single-copy core proteins using LG + CAT substitution model in individual human distal lung streptococci (red gradient) and closely related type strains (gray gradient). Bar charts show total number of GH family proteins (y-axis) in individual genomes (x-axis).

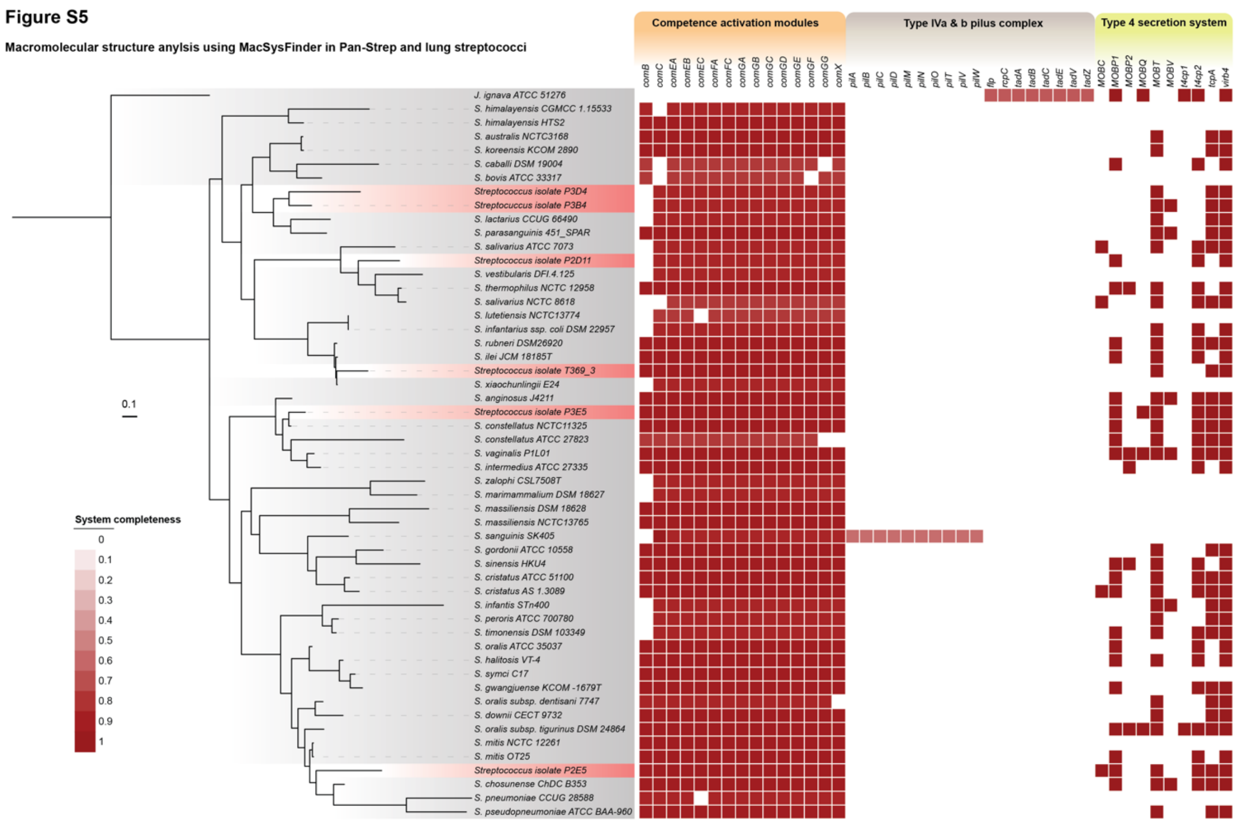


**Figure S5.** Heatmap showing completeness of macromolecular structures group into categories (colored gradient stripes) depicted according to maximum-likelihood phylogeny computed by FastTree on concatenated amino acid sequences of 315 single-copy core proteins using LG + CAT substitution model in individual human distal lung streptococci (red gradient) and closely related type strains (gray gradient).

### Figure S6

**Genomic arrangement and synteny of capsule genes**

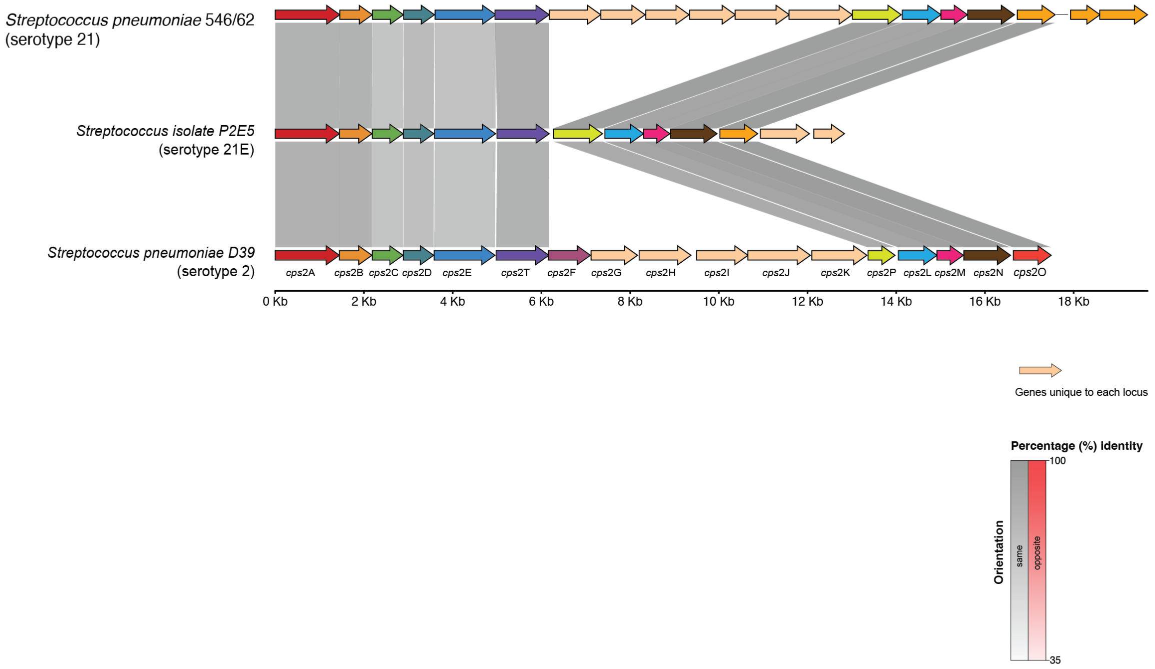


**Figure S6. Streptococcal capsular polysaccharide synthesis operon comparison and its relative genomic arrangement.** *cps* genes (colored arrows) in the type strain *S. pneumoniae* D39 (serotype 2), S. pneumoniae 546/62 (serotype 21) and *Streptococcus* isolate sp. nov. P2E5. Gene plot for genomic coordinates and synteny plots generated by pyGenomeViz comparing the percentage identity (shading depth) and relative gene orientation (grey for links with the same direction, pink for reverse direction).

**Figure S7**

**Phylogenetic comparison of human lung isolates with reference type strains and human oral isolates from expanded Human Oral Microbiome Database (eHOMD)**

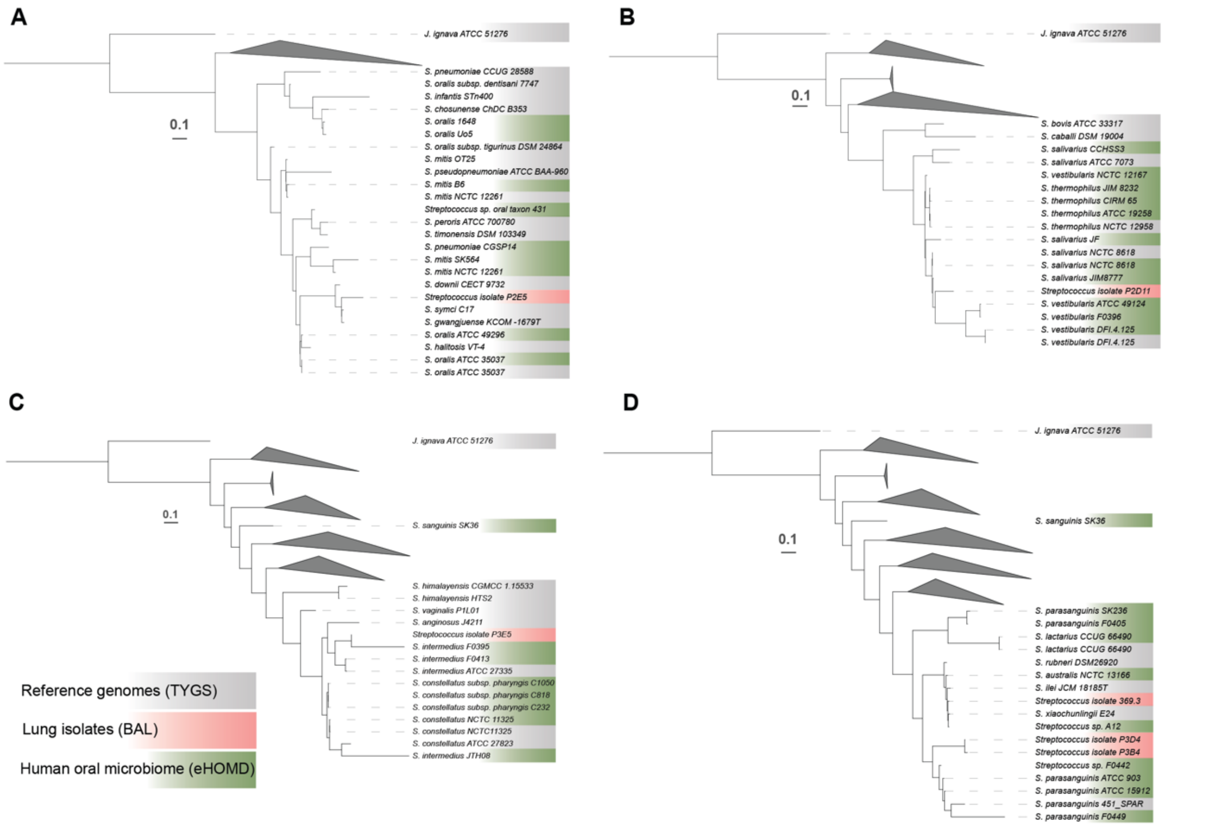


**Figure S7. (A-D) Phylogenetic comparison of individual distal lung streptococci with closely related human oral bacteria.** Single copy core-genome phylogeny comparing reference genomes from TYGS (gray gradient), human oral streptococci from eHOMD (green gradient) with individual lung streptococcal isolates cultivated from BAL (red gradient). Maximum-likelihood tree computed by FastTree on concatenated amino acid sequences of 26 single-copy core proteins using LG + CAT substitution model, collapsed to specifically focus on each isolate.

**Supplementary Table Legends**

**Table S1:** Result of the TYGS species identification showing Pairwise comparisons of user genomes vs. type-strain genomes. The dDDH values are provided along with their confidence intervals (C.I.) for the three different GBDP formulas formula $d_0$, $d_4$ and $d_6$ and G+C content difference (in %).

**Table S2:** Strains in this dataset automatically determined closest type strains by TYGS. Listed are NCBI Assembly accession with corresponding strain names.

**Table S3:** Identification and antibiotic test summary table containing results of hemolysis, optochin test, serotyping and MALDI-TOF and antibiotic susceptibility of six human distal lung streptococci.

**Table S4:** Results of Streptococcal identification using API® 20 Strep panel (Biomérieux, France) containing a summary and different result sheets for individual human distal lung *Streptococcus* generated by apiweb™.

**Table S5: Summary table of** Clusters of Orthologous Groups (COG) analysis in individual isolates and shared genes using eggNOG mapper.

**Table S6:** Output table from a custom rule-based pipeline for predicting metabolic pathways and macromolecular structure with completeness scores.

**Table S7:** Output summary of antibiotic resistance genes found in all genomes using ABRIcate tool.

**Table S8:** Output summary of virulence factors found in all genomes using ABRIcate tool.

**Table S9:** Closest strains to six lung isolates determined by BLASTN against eHOMD along with TYGS reference genomes. Listed are NCBI Assembly accession with corresponding strain names.

**Supplementary Datasets**

All datasets were submitted to Zenodo: https://doi.org/10.5281/zenodo.10220079

**Dataset S1 Lung_Streptococcus_genomes_metaQUAST:** MetaQUAST (Quality Assessment Tool for Metagenome Assemblies) output after genome sequencing and assembly of six lung streptococci. This includes HTML and PDF reports, summary statistics including total contigs, assembly size, and N50. Coverage analysis assesses how well reference genomes are represented, contig length distribution plots visualize contig length ranges, mis-assembly analysis detects potential errors and graphical representations to visualize assemblies.

**Dataset S2 Lung_streptococcus_isolate_genomes:** Nucleotide FASTA files of 6 lung streptococcal isolates obtained that were obtained via whole genome sequencing.

**Dataset S3 Lung_isolates_genome_annotation_prokka:** Output folders after annotation of six lung streptococcal isolates with PROKKA. This includes protein FASTA, GenBank files and GFF annotations.

**Dataset S4 TYGS_dDDH_analysis:** Contains results of TYGS analysis from DSMZ including downloadable reports. Outputs including taxonomic identification with genus, species, and strain details, a TYGS index for tracking genomes, genome quality assessment metrics, GBDP whole genome and 16S rRNA phylogenetic tree files, comparisons with reference type strains in the TYGS database with table.

**Dataset S5 Reference_type_strains_TYGS_genomes:** Nucleotide FASTA files of 47 closely related reference *Streptococcus* genomes listed by TYGS and downloaded from NCBI.

**Dataset S6 Reference_type_strains_TYGS_proteins:** Protein FASTA files of 47 closely related reference *Streptococcus* genomes listed by TYGS and downloaded from NCBI.

**Dataset S7 Lung_streptococcus_isolate_proteins:** Protein FASTA files of 6 six lung streptococcal isolates.

**Dataset S8 OrthoFinder_core_genome:** OrthoFinder is a bioinformatics tool that offers comprehensive outputs for orthology inference across multiple genomes. The output includes overall statistics, gene duplication information, orthologous genes, orthologous gene tree, single copy orthologous genes and STAG evolutionary trees.

**Dataset S9 Pan-Strep_BLAST_db:** BLAST database using the *makeblastdb* command of NCBI datasets command line tool. This is constructed using Protein

FASTA files of 47 closely related reference *Streptococcus* genomes listed by TYGS and downloaded from NCBI.

**Dataset S10 OrthoVenn_cluster_files:** OrthoVenn is a web-based tool for orthologous gene comparison. Downloadable results include Venn diagrams depicting shared and unique orthologous clusters amongst species, tabular results detailing genes within each cluster and their annotations. Functional enrichment analysis for Gene Ontology terms and KEGG pathways are also provided. enhances biological insights.

**Dataset S11_COG_analysis:** Results of COG analysis of six lung streptococcal isolates individually using eggNOG (evolutionary genealogy of genes: Non-supervised Orthologous Groups) webtool. The output includes information on Clusters of Orthologous Groups (COGs) categorizing them into functional groups such as metabolism, information storage and processing, and cellular processes and signalling.

**Dataset S12_CAZymes_lung_streptococci:** Results of CAZyme analysis using a custom rule-based pipeline mostly based on dbCAN (Database for Carbohydrate-Active enZymes) provides information on the carbohydrate-active enzymes present in genomic datasets. The output includes the annotation of enzymes involved in the degradation, modification, or biosynthesis of carbohydrates: glycoside hydrolases (GH), glycosyltransferases (GT), carbohydrate-binding modules (CBM), Auxillary Activities (AA), Carbohydrates Esterases (CE) and Polysaccharide lyases (PL).

**Dataset S13_pneumolysin_analysis:** Results alignment and phylogeny of Pnuemolysin protein in *Streptococcus pneumoniae*, *Streptococcus pseudopneumoniae* and Streptococcus isolate P2E5 found by ABRIcate analysis. Visual plots by pyGenomeViz.

**Dataset S14 Dataset S14_capsule_analysis:** Results from BLAST analysis of *Streptococcus pneumoniae* D39 capsular biosynthesis operon genes against the Pan-Strep (Dataset S10). Extracted of matching genes followed alignment and phylogeny. Visual plots by pyGenomeViz.

**Dataset S15_Lung_isolate_HOMD_TYGS_comparison:** Protein FASTA files of 47 closely related reference *Streptococcus* genomes listed by TYGS and downloaded from NCBI, 6 six lung streptococcal isolates and 47 streptococcal genomes from downloaded from human oral microbiome database (eHOMD).
